# Chromatin context-dependent deacetylation by the asymmetric Rpd3L

**DOI:** 10.64898/2026.01.08.698523

**Authors:** Heyu Zhao, Huadong Li, Chi Wang, Xuechen Yang, He Li, Binqian Zou, Shuqi Dong, Nan Zhang, Yuxing Zhou, Li Yi, Ying Zhang, Yixuan Xie, Dajiang Qin, William Chong Hang Chao, Duanqing Pei, Jun He

## Abstract

The regulation of gene expression requires precise control of chromatin-associated complexes that respond to diverse structural and epigenetic cues. The Rpd3 Large (Rpd3L) complex is a Sin3 histone deacetylase (HDAC) complex that dynamically adapt to chromatin states to reinforce transcriptional silencing, yet the mechanisms governing the catalytic activation in chromatin context-dependent manner remain unclear. Here we present the cryo-EM structure of Rpd3L bound to both mono- and di-nucleosome substrate at near-atomic resolution, uncovering a substrate-guided allosteric activation mechanism. Rpd3L adopts an asymmetric architecture, in which the proximal catalytic module anchors the first nucleosome, while the Sin3 PAH domains engage linker DNA to reposition a second nucleosome. This spatial configuration brings the distal catalytic module into proximity with chromatin and unlocks its latent deacetylase activity. Biochemical and mass spectrometry analyses confirm that dual nucleosome engagement selectively enhances Rpd3L activity and broadens substrate specificity. Together, these findings establish a hierarchical mechanism by which Rpd3L interprets histone modifications and nucleosome organization to modulate its enzymatic output at promoter regions. Our study provides a framework for understanding higher-order chromatin repression mechanisms by chromatin-regulation complexes and co-repressors.

## Introduction

Histone deacetylases complexes (HDACs) play central roles in establishing repressive chromatin states by removing acetyl groups from histone tails, thereby regulating gene expression and genome stability(1–5). Among these, Sin3 HDACs form conserved multi-subunit complexes from yeast to human(6–9), functioning in transcriptional repression(10–13) and contributing to key cellular processes such as differentiation and development(14–16). In *Saccharomyces cerevisiae*, the sole Sin3 isoform partners with the catalytic subunit Rpd3 to form two distinct complexes—Rpd3L and Rpd3S—which differ in subunit composition and chromatin targeting (17–19). Rpd3S deacetylates histones within gene bodies in a H3K36me3-dependent manner to suppress cryptic transcription(18,20,21), whereas Rpd3L preferentially targets H3K4me3-enriched promoter regions (10,22,23), reinforcing transcriptional silencing and genome integrity (24).

Recent studies reported that Rpd3L features a unique asymmetric architecture, with two Rpd3 catalytic modules arranged around a T-shaped supporting module and one Rpd3 subunit is autoinhibited by the Rxt2 subunit(25–27). This asymmetry suggests that Rpd3L may employ a distinct, modular mode of regulation like Rpd3S, functionally divergent from other class I HDACs(28,29). In addition to its structural asymmetry, the functional complexity of Rpd3L complex is also highlighted by its ability to interact with various co-repressors(30–33), integrating structural plasticity with chromatin context-dependent enzymatic activity, positioning it as a key player in hierarchical chromatin engagement and transcriptional repression regulation(9,34).

Despite this, the molecular basis by which Rpd3L engages nucleosomal substrates and coordinates its dual catalytic modules remains poorly understood. Here, using single-particle cryo-electron microscopy (cryo-EM) and biochemical assays, we set out to investigate how Rpd3L’s asymmetric modules orchestrate nucleosome recognition and deacetylation in a chromatin context-dependent manner. By using mono-nucleosome template, we provided the structural insights of nucleosome recognition by Rpd3L, implying how Rpd3L initiates the engagement on chromatin via its proximal module. We further show that Sin3 PAH domains, together with a second nucleosome, serve as an allosteric activator of Rpd3L’s distal module, unlocking its enzymatic potential. Through mutagenesis assays, we provided evidence that multi-nucleosome engagement is required to fully activate Rpd3L, providing direct evidence for its trans-acting regulation. Taken together, our findings uncover a previously unrecognized mechanism in which Rpd3L’s asymmetric catalytic modules are differentially regulated by nucleosome topology and positioning, allowing it to interpret and respond to chromatin architecture to achieve locus-specific transcriptional repression.

## Methods

### Protein expression and purification

The component subunits of *S. cerevisiae* wild-type Rpd3L (Rpd3, Ume1, Sap30, Cti6, Rxt3, Sin3(214–1536), Dep1, Rxt2 and Pho23) were amplified respectively from yeast genomic DNA by PCR and were cloned into modified MultiBac vectors for baculovirus expression. The double StrepII tag with cleavable TEV site was engineered at the N-terminal of Sin3(214–1536). Based on wild-type Rpd3L, mutant Rpd3L MultiBac vectors were constructed: one lacking the entire Rxt3 subunit (Rpd3L^ΔRxt3^), and another containing a truncated Sin3 fragment lacking the PAH1 and PAH2 domains (Rpd3L(Sin3 ^661–end^)). Baculoviruses containing all the component subunits of Rpd3L were mixed in a proper ratio to infect High Five insect cells to co-express the Rpd3L complex for 54 h. Cells were harvested and suspended in buffer A (50 mM HEPES pH 7.5, 350 mM NaCl, 0.5 mM TCEP, 10% glycerol) supplied with 1 mM PMSF, protease inhibitor cocktail, and supernuclease. Clarified cell lysate was loaded onto a streptavidin column for affinity purification. After using the buffer A with 2.5 mM Desthiobiotin to elute the protein complex, the fractions containing the target complex were diluted to buffer B (50 mM HEPES pH 7.5, 80 mM NaCl, 0.5 mM TCEP, 10% glycerol) and then be purified by the Resource Q column (Cytiva), and eluted with a gradient 0%-100% buffer C (50 mM HEPES pH 7.5, 1M NaCl, 0.5 mM TCEP, 10% glycerol). Finally, the protein complex was further purified by gel filtration in buffer D (50 mM HEPES pH 7.5, 300 mM NaCl, and 0.5 mM TCEP).

### Preparation and purification of histone octamer

The pETDuet-1- *Xenopus laevis* vector was modified to enable polycistronic expression of histone octamer(35). *Escherichia coli* Rosetta (DE3) cells harboring this vector were used. The collected bacterial pellets were lysed in buffer (20 mM Tris pH 8.0, 500 mM NaCl, 0.1 mM EDTA, 0.5 mM TCEP), and the supernatant was applied to a Heparin affinity column (Cytiva). After washing with 500 mM NaCl buffer, bound proteins were eluted using a salt gradient from 500 mM to 2 M NaCl. To eliminate DNA contamination, size-exclusion chromatography was performed in a high-salt buffer (20 mM Tris pH 8.0, 2 M NaCl, 0.5 mM EDTA, 0.5 mM TCEP). In order to mimic the H3K4me3 modification, histone H3 C110 was mutated to alanine and H3K4 was mutated to cysteine to enable site-specific installation of methyl-lysine analogs (MLAs). Histone octamers containing these double mutations were expressed and purified using the same procedure as described for the wild-type histone octamer. Site-specific installation of the methyl-lysine analog at H3K4 was performed as previously described (36), using (2-bromoethyl) trimethylammonium bromide to generate H3K4me3 (MLA)-containing histone octamers. Endogenous histone octamers were extracted from 293F cells using the same protocol used for wild-type histone octamer purification.

### Nucleosome reconstitution

DNA templates for nucleosome assembly were generated by PCR amplification using the Widom ‘601’ positioning sequence. Mono-nucleosomes were assembled using ‘601’ DNA fragments with 20 bp flanking linker DNA on both ends (20N20). Di-nucleosomes with varying inter-nucleosomal spacings were assembled using synthetic DNA templates containing two ‘601’ sequences separated by linker regions of 20 bp (30N20N10), 28 bp (30N28N10), 32 bp (30N32N10), 40 bp (30N40N10), or 55 bp (30N55N10). PCR products were purified by ion exchange column. Peak fractions were collected and subsequently precipitated with isopropanol. For fluorescence-based assays, Cy5 and FAM fluorophores were incorporated into the reverse primers of the 20N20 and 30N32N10 DNA fragments, respectively. Nucleosomes were reconstituted using the ‘double bag’ dialysis method(37). DNA and histone octamers (1:1.1 molar ratio) were mixed in high-salt buffer (20mM Tris-HCl, pH 8.0, 2M NaCl) and placed in dialysis buttons inside a dialysis bag containing 50 ml of the same buffer. The bag was dialyzed overnight at 4 °C against 1L of 1M NaCl buffer, then transferred to 1 L of low-salt buffer (50 mM NaCl) for 5-6 h. Finally, the dialysis buttons were dialyzed separately in fresh low-salt buffer for 3-4 h. Reconstituted nucleosomes were collected and analyzed before use.

### Sample preparation and cryo-EM data collection

The Rpd3L-mono-nucleosome complex was prepared by mixing Rpd3L with nucleosome(20N20) at a molar ratio of 0.9:1, while the Rpd3L-di-nucleosome complex was obtained by mixing Rpd3L with di-nucleosome (30N32N10) at a ratio of 0.5:1. To stabilize the complexes, we employed the GraFix method(38). A 6 ml top gradient solution (50 mM NaCl, 20 mM HEPES pH 7.5, and 10% glycerol) was layered over a 6 ml bottom solution (50 mM NaCl, 20 mM HEPES pH 7.5, 30% glycerol, and 0.125% glutaraldehyde) in a tube (Beckman). Gradients were formed using a Gradient Master (BioComp). For the Rpd3L-mono-nucleosome complex, 170 μl of sample was loaded and ultracentrifuged at 4 °C for 14 h at 35,000 rpm (Beckman SW-41Ti rotor). For the Rpd3L-di-nucleosome complex, 150 μl of sample was ultracentrifuged under the same conditions for 12 h at 32,000 rpm. Fractions were collected, and the optimal ones were dialyzed in buffer (20 mM HEPES, pH 7.5, and 50 mM NaCl) before cryo-EM grid preparation.

Cryo-EM grids were prepared by applying 3 μl of the sample onto glow-discharged Quantifoil Cu R1.2/1.3 300-mesh grids. The grids were blotted for 4 s and plunge-frozen in liquid ethane using a Mark IV Vitrobot (FEI). Micrographs were collected on a Titan Krios G4i (Thermo Fisher Scientific) operating at 300 kV. A total of 11,370 images for the Rpd3L-mono-nucleosome complex and 11,135 images for the Rpd3L-di-nucleosome complex were recorded using a Falcon4 direct electron detector equipped with a SelectrisX energy filter (10 eV slit width). Each image was acquired using EPU software at a nominal magnification of 165,000×, corresponding to a pixel size of 0.71 Å/pix. The defocus range was between −0.8 and −2.4 µm, with an accumulated total dose of approximately 50 e^−^/Å^2^.

### Image processing

All EER movies were processed using CryoSPARC(39), with initial motion correction and CTF estimation performed in real time via CryoSPARC Live. For the Rpd3L-mono-NCP complex, 10,157 micrographs were manually selected for further processing. Two separate batches of particles were automatically picked using blob picking, with the second batch selected by excluding the final set of particles from the first batch. The first batch of 1,159,655 particles was extracted with a 720-pixel box (binned 4×) and subjected to 2D classification, retaining 543,625 particles for ab initio reconstruction. Class 6 from this reconstruction was selected as a template for Topaz particle picking (600-pixel box, binned 4×). The Topaz-picked particles were then processed in the same manner, and Class 5 was selected as an additional template. Particles from Class 6 (first batch) and Class 5 (second batch) were re-extracted at 600 pixels, bin-1, and high-quality particles were merged and duplicates removed, yielding 248,172 bin-1 particles. The second batch underwent parallel multi-reference guided 3D classification performed twice. Particles were initially extracted with a 600-pixel box (binned 4×) for the classifications. After each round, duplicate particles were removed and the remaining particles merged before repeating the procedure. The final set of 349,786 particles was re-extracted at 600 pixels, bin-1. After removing duplicate particles, merging with the first dataset yielded a total of 451,544 particles for homogeneous refinement. The volume was split into three and subjected to local refinement. The local volume maps were combined in the model building.

For the Rpd3L-di-NCP complex, 1,940 micrographs were manually selected from an initial set of 2,020 micrographs for blob picking and 2D classification. Particles were extracted with a 600-pixel box size (binned 4×), resulting in 169,426 particles for *ab initio* reconstruction. Class 5 was chosen as the initial model for further processing. A second dataset containing 9,115 micrographs was collected, of which 9,085 were selected for additional rounds of particle picking. To improve the density of nucleosome 2, particles were divided into two groups based on defocus parameters. After 2D classification and refinements, 205,066 particles (re-extracted at bin-1) were selected for local refinement of the Rpd3L, nucleosome 1, nucleosome 2, and linker DNA, resulting in local density maps with the following resolutions: Rpd3L at 3.8 Å, nucleosome 1 at 2.9 Å, nucleosome 2 at 3.6 Å, and linker DNA at 5.9 Å. The local volume maps were combined for the model building.

### Model building

The nucleosome structure (PDB ID: cryopros-uniform-pose) (40) and the Rpd3L complex (PDB ID: 8HPO) (25) were used as initial structural templates. The model was docked into the cryo-EM density maps using UCSF ChimeraX(41). AlphaFold3(42) was employed to predict the structures of PAH1 and PAH2 domains of Sin3. The initial fitted models were manually adjusted in COOT(43), followed by multiple rounds of real-space refinement using Phenix(44) and Namdinator(45). The structure figures were generated using PyMOL (https://pymol.org/2/) or UCSF ChimeraX.

### Cross-linking mass spectrometry

Rpd3L was incubated with nucleosomes at a molar ratio of 0.9:1 in the presence of 2 mM BS3 (bis(sulfosuccinimidyl)suberate) on ice for 1 h. The cross-linking reaction was quenched with 50 mM Tris buffer pH 8.0, maintaining the same pH as the HEPES buffer. The sample was then flash-frozen in liquid nitrogen and stored at −80 °C. The cross-linked complex was enzymatically digested and pre-fractionated using HPLC. Following desalting, the sample was analyzed by LC-MS/MS. The resulting data were processed using pLink2 software(46) and visualized with xiView(47).

### EMSA and nucleosome competition assay

Using the previously described fluorescently labeled nucleosome method, 5’-FAM was labeled on di-nucleosome, and Cy5 was labeled on mono-nucleosome. Rpd3L and the two nucleosomes were first incubated at a ratio of 2:1 (pmol) for 30 min. Subsequently, either mono-nucleosome or di-nucleosome was added to the reaction mixture (10 μl), followed by an additional 30 min incubation. The concentration gradients were as follows: 0, 0.15, 0.3, 0.45, 0.6, 0.75, 0.9, 1.05, 1.2, 1.35 and 1.5 pmol (di-nucleosome); 0, 0.3, 0.6, 0.9, 1.2, 1.5, 1.8, 2.1, 2.4, 2.7 and 3 pmol (mono-nucleosome). The samples were separated using 4.5% native gel. The two fluorescent signals were detected using the Invitrogen iBright 1500 imaging system (Thermo Fisher Scientific), with excitation/emission settings of 495/520 nm for FAM and 640/665 nm for Cy5, respectively.

### Fluorescence polarization assay

FAM was incorporated into mono-nucleosome and di-nucleosome as described above. Binding affinities were measured by incubating 20 nM of each 5’-FAM-labeled nucleosome with a two-fold serial dilution of Rpd3L components in binding buffer (20 mM Tris pH 7.5, 100 mM NaCl, 1 mM EDTA, 2 mM DTT, 5% glycerol, 0.01% NP-40, 0.01% CHAPS, 0.1 mg/ml BSA). For the wild-type Rpd3L complexes, protein concentrations ranged from 2 μM to 0.000244 μM. For Rpd3L(Sin3 661–end), protein concentrations ranged from 1 μM to 0.000244 μM. After a 30 min-incubation at room temperature, 40 μl of each reaction was transferred to a 384-well low-volume black, round-bottom, non-binding surface microplate (Corning). Fluorescence polarization was read on a BioTek Synergy Neo2 plate reader equipped with excitation and emission filters (485/20 nm and 528/20 nm, respectively) and a polarizer; the gain was set to 50 for all measurements. Unless otherwise noted, each experiment was performed in triplicate and the mP values from two independent scans were averaged. Background subtraction and data normalization were carried out as previously described, and dissociation constants (Kd) were obtained by fitting to a specific-binding with Hill-slope model in GraphPad Prism ( http://www.graphpad.com).

### Boc-Lys(Ac)-AMC deacetylation assay

Boc-Lys(Ac)-AMC is a fluorogenic substrate for histone deacetylases(48). HDAC deacetylates the substrate and the product (Boc-Lys-AMC) is treated with trypsin releasing fluorescent AMC. In a 20 µl reaction mixture, nucleosomes at 100 nM and 37.5 µM Boc-Lys(Ac)-AMC were included, with Rpd3L^WT^ at 0, 5, 10, 15, and 20 nM. The concentrations of Rpd3L^ΔRxt3^ were 0, 1, 2, 3, and 4 nM. The working buffer consisted of 100 mM NaCl, 20 mM HEPES pH 7.5, 5% glycerol, 0.5 mM TCEP, and 1 µM ZnSO₄. After a 20 min reaction, deacetylation was terminated using 10 µM trichostatin A (TSA). Finally, 10 µl of 2 mg/ml trypsin was added for digestion for 30 min before fluorescence signal detection. The data were analyzed using the Simple Linear Regression method in GraphPad Prism to determine the reaction rate relationships of different components.

### Histone Deacetylation Assay

To evaluate the deacetylation activity of the Rpd3L complex, nucleosomes were reconstituted using endogenously acetylated histone octamers. Mono-nucleosomes were assembled with 147 bp Widom 601 DNA flanked by 20 bp linker sequences on both ends (20N20). Di-nucleosomes were assembled using DNA fragments containing two 601 positioning sequences separated by a 32 bp linker region (30N32N10), generating variable internucleosomal spacing. Deacetylation reactions were performed in a total volume of 10 μL. Mono-nucleosomes were used at a final concentration of 250 nM, and di-nucleosomes at 125 nM. Rpd3L was titrated at final concentrations of 0, 1, 2, 4, 8, and 16 nM. Reactions were carried out at 25 °C for 30 minutes and quenched by the addition of SDS sample loading buffer, followed by heating at 98 °C for 5 minutes. Proteins were resolved by 15% SDS-PAGE, transferred to PVDF membranes, and subjected to Western blotting. Acetylation levels were probed using site-specific antibodies against H3K9ac, H3K18ac, H3K27ac, H4K5ac, H4K12ac, and H2BK12ac. Antibodies against total H3 and H4 were used as loading controls. Signals were detected using enhanced chemiluminescence (ECL).

### LC-MS/MS Analysis

Samples were analyzed on an ACQUITY UPLC M-Class system (Waters) coupled to a ZenoTOF 7600 mass spectrometer (SCIEX). An OptiFlow TurboV ion source equipped with a microflow probe was utilized. Peptides were separated using a nanoEase M/Z Peptide BEH C18 column (2.6 μm, 0.3×150 mm, Waters) at a constant flow rate of 5 μl/min with mobile phase buffers A and B (buffer B: ACN containing 0.1% FA, v/v). The chromatography gradient consisted of: 0-2 min, 3% B; 2-12 min, 3-32% B; 12-12.5 min, 32-80% B; 12.5-14 min, 80% B; 14-14.5 min, 80-3% B; 14.5-18 min, 3% B. Ion source Gas 1, Gas 2, and curtain gas were set to 20, 60, and 35 psi, respectively. The mass spectrometer was operated in positive ZenoSWATH DIA mode with collision-induced dissociation (CID) fragmentation. The TOF MS scan range was m/z 120-1400 with a 0.1 s accumulation time. Dynamic collision energy was enabled and ranged from 14 to 46 V. Product ions were monitored in the range m/z 120-1200 with a 10 ms accumulation time. WIFF data files were converted using EpiConverter and analyzed with EpiProfile and Skyline software.

## Results

### Rpd3L engages the mono-nucleosome asymmetrically through the proximal module

To visualize the structural architecture of the nucleosome bound Rpd3L complex, we recombinantly expressed and purified *S. cerevisiae* Rpd3L with subunits Sin3, Rpd3, Ume1, Sds3, Dep1, Pho23, Rxt2, Sap30, Rxt3, Cti6 in insect cells using a baculovirus expression system (Fig. 1A, Supplementary Fig. S1A and B)(49). We reconstituted a nucleosome core particle (NCP) with two extra 20 bp DNA linkers flanking the Widom 601 sequence (187bp) and then prepared Rpd3L–mono-nucleosome complexes for cryo-EM studies(50). The resulting high-resolution reconstruction map, which resolved the nucleosome and the Rpd3L complex at 2.7 Å and 2.9 Å resolution respectively, enabled us to model nearly all subunits of the purified Rpd3L complex, with the exception of Ume1, for which no corresponding density was observed (Fig. 1B and D, Table 1, Supplementary Fig. S2 and 3).

**Fig. 1.**
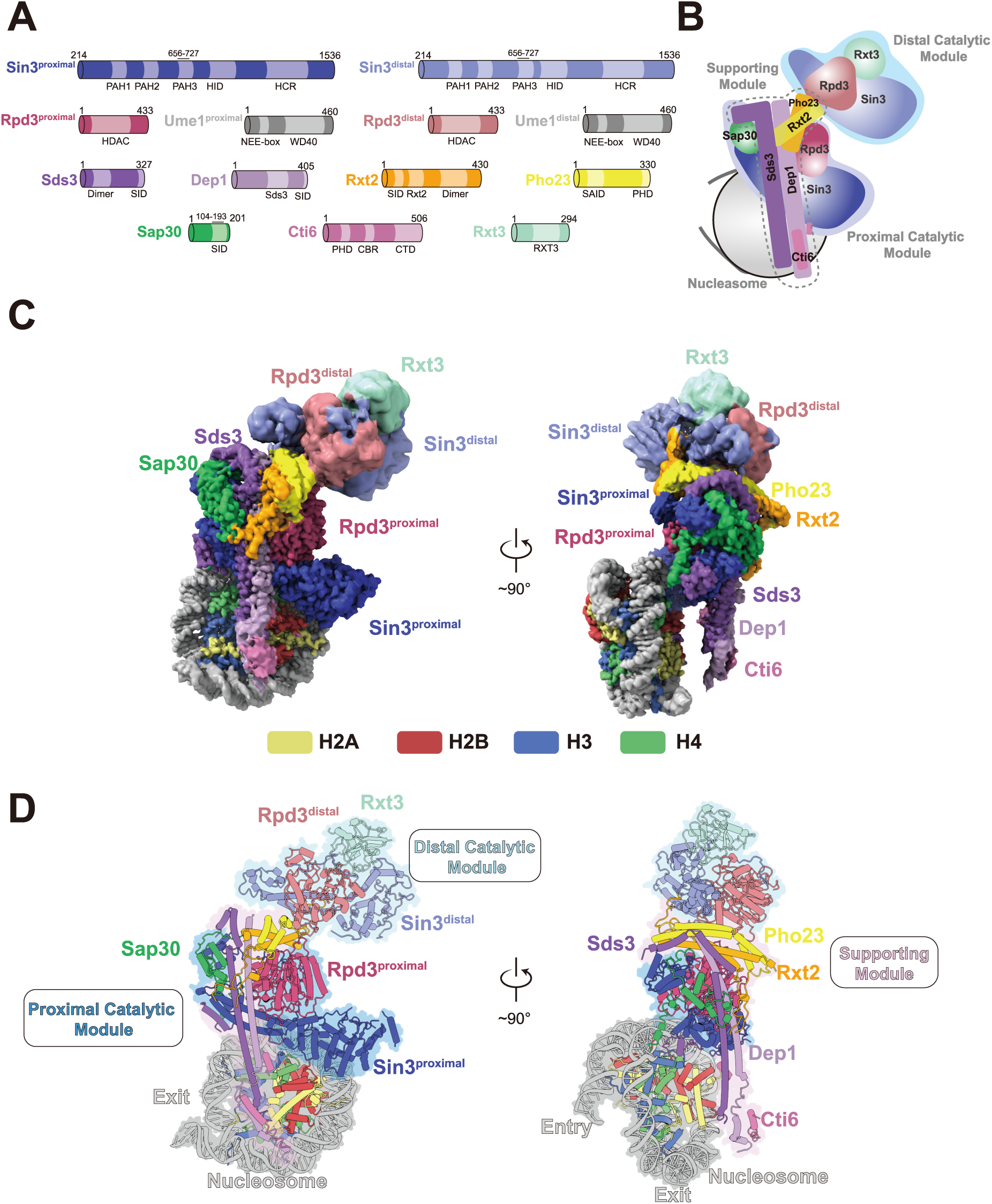
Overall structure of the Rpd3L bound to the mono-nucleosome. **A**, Domain organization of Rpd3L subunits. **B**, Schematic representation of the Rpd3L bound to the mono-nucleosome. The distal and proximal catalytic domains are highlighted with light blue and blue backgrounds, respectively, while the T-shaped supporting module is outlined with dashed lines. **C**, Top view (left) and side view (right) of the cryo-EM composite map of the Rpd3L bound to the mono-NCP. **D**, Corresponding atomic model in cylindrical view. The distal and proximal catalytic modules are highlighted with light blue and blue backgrounds, respectively, while the T-shaped supporting module is highlighted in light pink. In all panels, subunits are colored according to the scheme shown in A.

The Rpd3L complex adopts an intricate and asymmetric architecture upon binding to a mono-nucleosome substrate, closely resembling its apo form(25,26). The nucleosome interacts asymmetrically with the complex, engaging primarily with one side. Structurally, Rpd3L consists of three distinct modules: the proximal catalytic module, which contains a Sin3-Rpd3 core pair along with co-regulatory components Sap30 and Cti6, that together directly interfaces with the nucleosome (Fig. 1D, left); the T-shape cross-brace supporting module primarily formed by two coiled-coil pairs of Sds3-Dep1 and Pho23-Rxt2, which together provides structural stability (Fig. 1D, right); and the distal catalytic module, which comprises the second Sin3-Rpd3 core pair associated factors Rxt3 (Fig. 1D, left). This organization underscores a functionally asymmetric mechanism of nucleosome recognition and histone deacetylation by Rpd3L (Fig. 1B).

Our structure reveals that Rpd3L engages the mono-nucleosome substrate primarily via its proximal module, establishing multivalent contacts. A key interaction involves the highly conserved polybasic surface of Sin3^proximal^ (K940, K941, K946, K1251), which forms hydrogen bonds and long-range electrostatic interactions with nucleosomal DNA at superhelix location (SHL) +3 (Fig. 2A–C, Supplementary Fig. S4). Basic anchor-mediated recognition of the H2A–H2B acidic patch is a common feature among chromatin regulators(51). We observed a defined side-chain density at the acidic patch, though the associated main-chain density was heterogeneous (Fig. 2E). Sequence analysis identified a lysine-and arginine-rich region in Cti6 (residues 370–430; Fig. 2D), and cross-linking mass spectrometry confirmed its interaction with the acidic patch (Supplementary Fig. S1C-E). These findings suggest that Cti6 engages the nucleosome acidic patch via any one of these lysine or arginine residues, leading to variability in backbone density positioning, as also suggested in previous work (26). Supporting this interpretation, we identified a previously unassigned density near the patch that fits Cti6 residues 244–252 (Fig. 2F), indicating that this intrinsically disordered segment in apo form becomes partially stabilized upon nucleosome binding. Additional electrostatic interactions were observed between nucleosomal DNA and the N-terminal helices of Sap30 and Sds3 (Fig. 1D, 2B, Supplementary Fig. S5A and B).

**Fig. 2.**
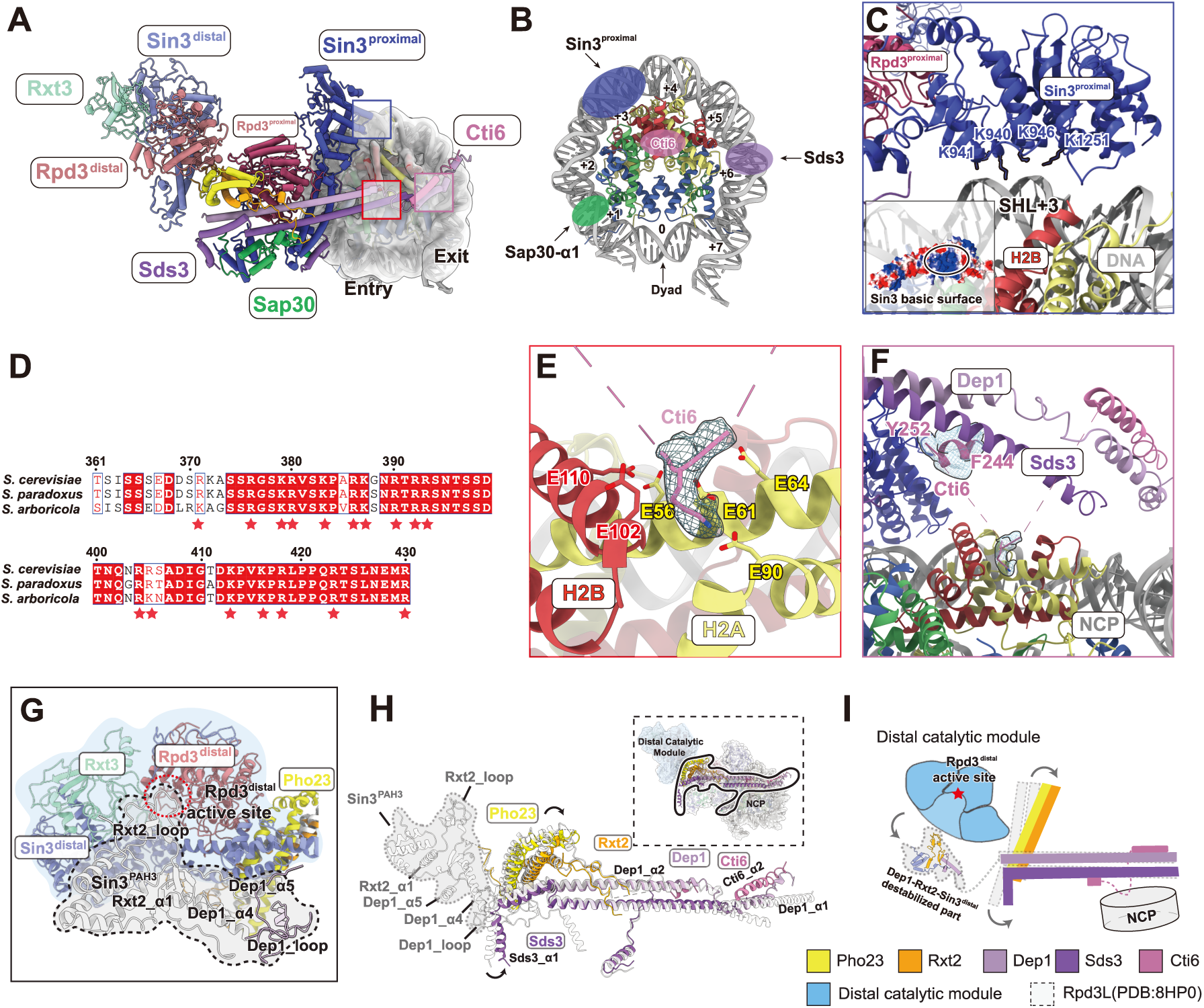
Details of Rpd3L complex engagement with the mono- nucleosome. **A**, Overview of the Rpd3L bound to the mono-NCP, highlighting the major interaction interfaces. The nucleosome density is shown as a cryo-EM map, filtered with a Gaussian low-pass filter. **B**, Schematic of the Rpd3L binding interfaces on the nucleosome. Nucleosomal DNA positions are labeled by superhelical locations (SHLs). **C**, Interaction of Sin3^proximal^ with nucleosomal DNA. The basic surface of Sin3^proximal^ interacts with nucleosomal DNA at SHL+3. Electrostatic surface potential (-/+5.0 kT/e) is mapped onto the Sin3^proximal^ surface to highlight charge distribution at the DNA-binding interface. **D**, Multiple sequence alignment of Cti6 orthologs. The displayed region corresponds to the central disordered segment located between modeled regions in Cti6. The alignment reveals a high enrichment of basic lysine and arginine residues (indicated by asterisks). **E**, Interaction of the lysine anchor of Cti6 with the H2A-H2B acidic patch. The local cryo-EM density corresponding to the lysine anchor is shown. **F**, Stabilization of a disordered Cti6 segment (F244-Y252) via interaction with the nucleosomal acidic patch. Local cryo-EM densities of both the segment and the lysine anchor are displayed. **G**, In the nucleosome-bound state, the Rxt2 loop (residues 69-88), along with Dep1_α4, Dep1_loop (purple), Dep1_α5, Rxt2_α1, and the PAH3 domain of Sin3^distal^, becomes flexible and is not resolved in the cryo-EM map. These regions are highlighted with dashed circles and shaded with a light grey background, representing the corresponding regions from the apo-Rpd3L model (PDB ID: 8HPO)(25).**H**, Structural comparison of the supporting module in apo Rpd3L (white) and mono-nucleosome-bound Rpd3L (colored), highlighting local conformational rearrangements upon nucleosome engagement. A global view of the complex is shown in the upper right corner, with the supporting module outlined in solid black to indicate its scaffold-like position within the entire assembly. Regions that become more flexible upon nucleosome binding, as inferred from the apo Rpd3L model, are outlined with dashed lines and shaded with a light grey background(25). **I**, Schematic illustration of nucleosome-induced conformational changes in the scaffold. Apo-state components are outlined in white with dashed lines; structural elements that become ordered or repositioned upon nucleosome binding are colored.

### Allosteric transition of Rpd3L distal catalytic module upon mono-nucleosome engagement

Within the Rpd3L complex, the proximal and distal catalytic cores (Rpd3^proximal^ and Rpd3^distal^) share highly similar structural features, but previous structural analyses of the apo form revealed distinct local conformations. Specifically, while the active site of Rpd3^proximal^ remains accessible, the catalytic site of Rpd3^distal^ is sterically occluded by a loop from Rxt2 (Fig. 2G grey, Rxt2_loop, Supplementary Fig. S5C and E), forming an auto-inhibitory arrangement, which is primarily maintained by interactions involving Dep1 α4-α5 (Fig. 2G, Supplementary Fig. S5C and S6) and the PAH3 domain of Sin3^distal^ (Fig. 2G grey, Sin3^PAH3^, Supplementary Fig. S5C and S6)(25,26). The evolutionary rationale for maintaining a second catalytically inactive HDAC core remains unclear.

Upon mono-nucleosome binding, we observed the disappearance of the Rxt2 inhibitory loop, accompanied by increased flexibility in surrounding supporting elements, including the Dep1 helices and the Sin3^distal^ PAH3 domain (Fig. 2G grey, Supplementary Fig. S5D). Comparison between apo and nucleosome-bound structures reveals that nucleosome engagement in the Rpd3L proximal module induces significant conformational changes in the T-shaped scaffold, which serves as a structural bridge between the proximal and distal modules (25,26). This engagement triggers a conformational shift that propagates along the Dep1-Sds3 coiled coil, leading to destabilization of the Dep1 and Sin3^distal^ domains (Fig. 2H, Supplementary Fig. S6). This structural rearrangement ultimately relieves Rxt2 auto-inhibition, making the Rpd3^distal^ catalytic site structurally accessible (Schematic Fig. 2I).

To confirm that Rpd3L’s deacetylase activity is enhanced upon nucleosome binding, we employed the fluorogenic substrate Boc-Lys(Ac)-AMC to quantitatively measure Rpd3L activity in the presence or absence of nucleosomes (48,52). Indeed, compared to the apo form, the nucleosome-bound Rpd3L complex exhibited significantly increased enzymatic activity at the same molar concentration, supporting that the allosteric conformational changes induced by mono-nucleosome binding effectively enhance Rpd3L’s catalytic activity (Fig. 3A).

**Fig. 3.**
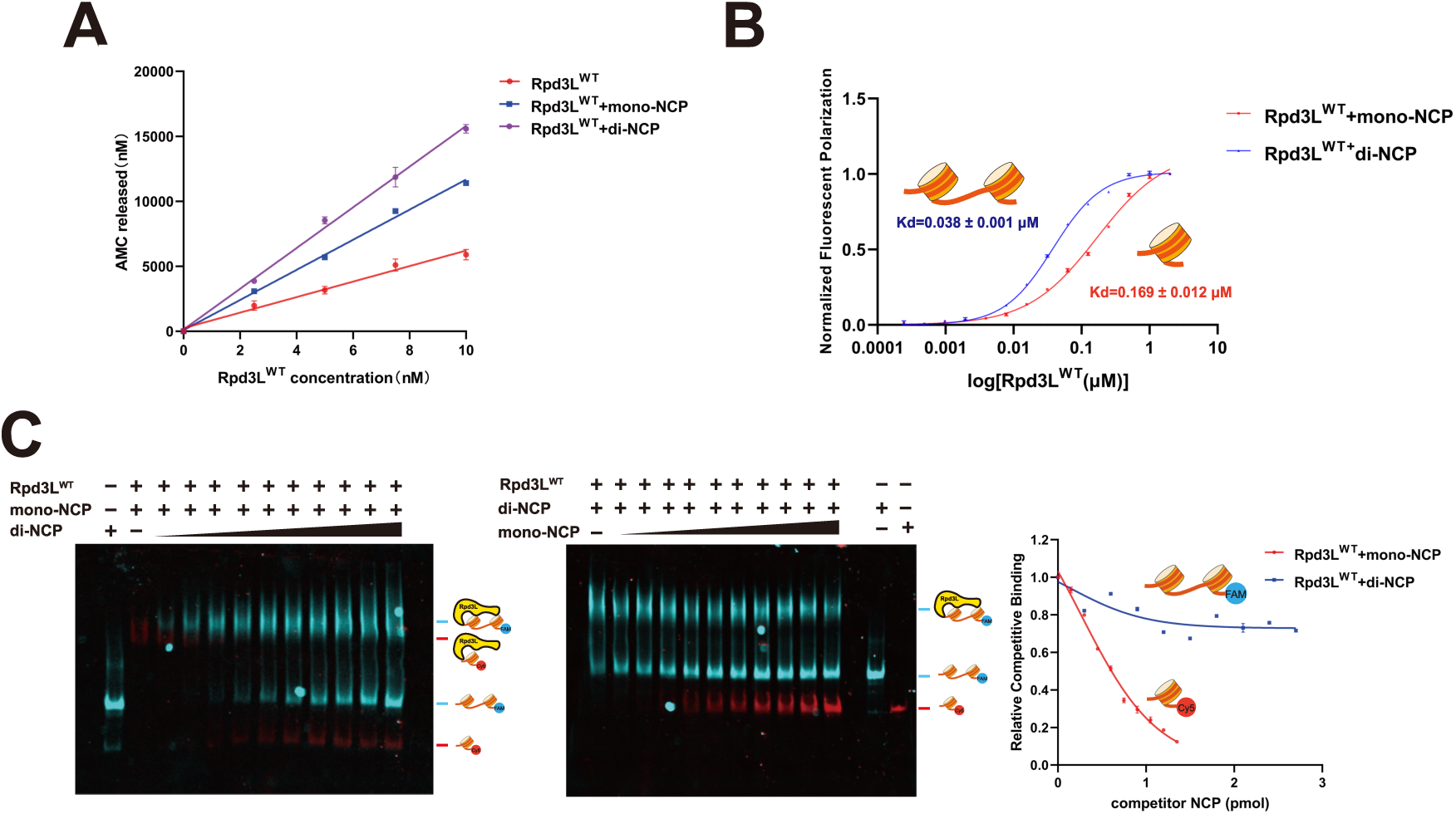
Rpd3L preferentially binds to di-nucleosomes over mono-nucleosomes. **A**, Deacetylation activity assay of Rpd3L in the presence of different substrates, including Rpd3L, Rpd3L + mono-NCP, and Rpd3L + di-NCP. The deacetylase activity of Rpd3L was measured using the fluorogenic substrate Boc-Lys(Ac)-AMC. Experiments were performed independently three times, and the error bars represent the variation between replicates (n = 3). **B**, Fluorescence polarization assay comparing wild-type Rpd3L binding to mono-NCP and di-NCP. Each measurement was performed in triplicate, with error bars showing the standard deviation (SD). The fitted Kd values are indicated, with the standard error (SE) calculated from the curve fitting procedure. **C**, Fluorescent images of the native gel, showing merged signals from both FAM and Cy5 channels. In Competition 1, di-NCP was titrated at 0, 0.15, 0.3, 0.45, 0.6, 0.75, 0.9, 1.05, 1.2, 1.35, and 1.5 pmol (left). In Competition 2, mono-NCP was titrated at 0, 0.3, 0.6, 0.9, 1.2, 1.5, 1.8, 2.1, 2.4, 2.7, and 3 pmol (middle). Quantitative analysis of fluorescence signals from the competition assays is shown on the right. The fluorescence intensity (grayscale) was measured in triplicate for each condition, and the values represent the average of these three measurements (n = 3).

Notably, increased enzymatic activity does not directly equate to full activation of Rpd3L distal module. While the Rpd3^distal^ catalytic site becomes structurally exposed, effective enzymatic activity on the nucleosomal substrate requires Sin3^distal^’s polybasic surface to be available for nucleosome binding. However, in the current conformation, this polybasic surface remains obstructed by Rxt3, preventing Sin3^distal^ from engaging the nucleosome (Supplementary Fig. S7). Thus, upon mono-nucleosome binding at the proximal module, the distal module transitions into a more flexible conformation with an accessible catalytic site but remains inactive to deacetylate nucleosomal substrate. This is consistent with our structural data, as we did not observe nucleosome engagement at the distal module in the complex structure. The implications of this structural state also prompted us to investigate Rpd3L in the context of a more physiologically relevant nucleosome template—the di-nucleosome.

### Rpd3L prefers di-nucleosome over mono-nucleosome as its substrate

Both Rpd3L and Rpd3S complexes have been proposed to bind di-nucleosome substrates. However, unlike Rpd3S, no direct studies have yet confirmed whether Rpd3L preferentially engages di-nucleosome substrates (53–55). Given that Rpd3L contains two catalytic modules, it has been hypothesized that Rpd3L also favors di-nucleosome templates for simultaneous catalytic reactions on separate nucleosomes (26). To test this hypothesis, we designed a di-nucleosome DNA template containing two 601 positioning sequences separated by a 32 bp linker DNA. We first employed fluorescence polarization assays to quantify the binding affinity of Rpd3L for nucleosome substrates. The results revealed that Rpd3L binds di-nucleosomes with approximately four-fold higher affinity than mono-nucleosomes (Fig. 3B).

To validate this binding preference of di-nucleosomes, we further conducted a fluorescence-based competition electrophoretic mobility shift assays (EMSA) (Supplementary Fig. S8G). To distinguish between mono-nucleosome and di-nucleosome substrates, we employed end-labeling with fluorophores FAM (blue) or Cy5 (red) (Supplementary Fig. S8A). The results indicated mono-nucleosomes were more readily displaced in binding reactions (Supplementary Fig. S8B-C, S8E). To account for the difference in nucleosome content, we repeated the assay with twice the amount of mono-nucleosome, yet Rpd3L remained preferentially bound to di-nucleosomes (Fig. 3C; Supplementary Fig. S8D). Furthermore, to determine whether linker DNA length contributes to this increased affinity, we tested di-nucleosomes with linker DNA ranging from 20 to 55 bp, and Rpd3L displayed consistent binding across all conditions (Supplementary Fig. S8F).

We also investigated whether Rpd3L exhibits differential enzymatic activity upon binding to di-nucleosome substrates. Using a deacetylation assay with the fluorogenic substrate Boc-Lys(Ac)-AMC, in which deacetylation by Rpd3L releases AMC that can be quantitatively monitored, we observed significantly enhanced deacetylation activity of Rpd3L in the presence of di-nucleosomes compared to its activity in apo form or when bound to mono-nucleosomes (Fig. 3A). Collectively, these findings provide strong evidence that Rpd3L preferentially engages di-nucleosome substrates by simultaneously contacting two nucleosomes and confirm that the enhanced catalytic potential of Rpd3L is triggered by di-nucleosome binding.

### Sin3 PAH1/2 domains direct a ‘guided landing’ architecture for Rpd3L engagement with di-nucleosome

To elucidate the molecular basis underlying Rpd3L’s preference for di-nucleosome substrates, we reconstituted di-nucleosomes *in vitro* using a 32-bp linker DNA. Cryo-EM analysis of the resulting Rpd3L-di-nucleosome complex yielded a high-resolution density map, clearly revealing two nucleosomes arranged at approximately a 100° angle, flanking the T-shaped scaffolding module and proximal catalytic module of Rpd3L; The distal module of Rpd3L, by comparison, displayed markedly increased flexibility relative to its conformation in the apo and mono-nucleosome-bound states (Fig. 4A, Supplementary Fig. S9 and 10, Table 1). In the resolved structure, the proximal catalytic module engages the first nucleosome (nucleosome 1) through a tripod-like arrangement, consistent with mono-nucleosome binding mode. The second nucleosome (nucleosome 2) is predominantly recognized by the N-terminal region of Sds3 and the PAH3 domain of Sin3^proximal^, both structurally bridged by the C-terminal region of Sap30 (Fig. 4A).

**Fig. 4.**
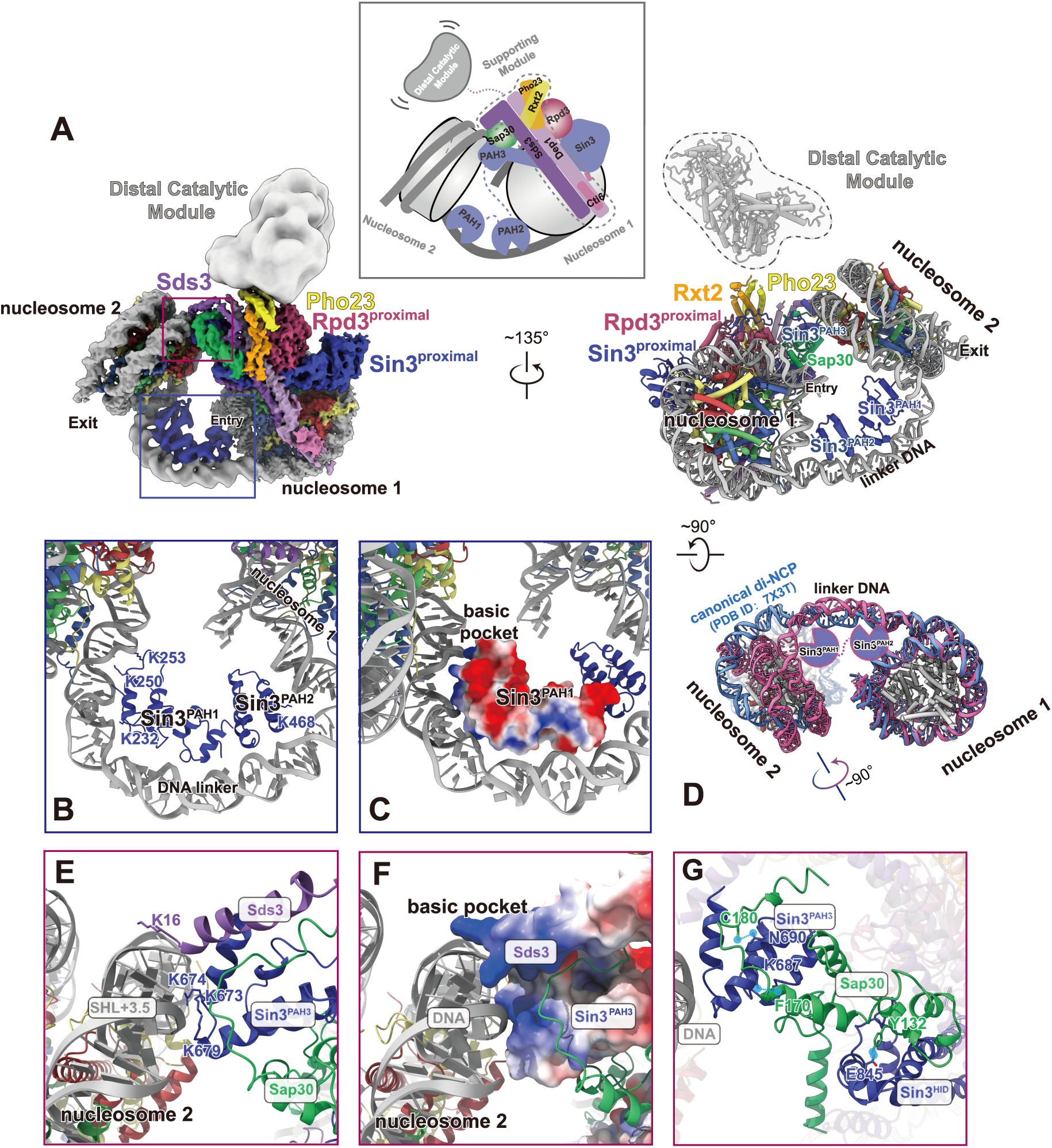
Details of Rpd3L complex bound with the di-nucleosome. **A**, Top (left) and side (right) views of the cryo-EM composite map and the cylindrical model of the Rpd3L-di-NCP complex. Subunits are colored according to the scheme shown in Fig.1A. The distal catalytic module of Rpd3L is highly flexible and thus appears fragmented in the density map; this region is indicated in grey. A Schematic illustration of the Rpd3L bound to di-NCP complex summarizes the multivalent interaction between Rpd3L and the di-nucleosome, with the distal catalytic module exhibiting structural flexibility. **B**, Enlarged view of the PAH1 and PAH2 domains of Sin3 in the proximal module, showing the interaction of the Sin3^PAH1^ and Sin3^PAH2^ domains with the linker DNA connecting the two nucleosomes. **C**, Electrostatic potential distribution of the PAH1 domain of Sin3^proximal^, shown in the same orientation as in B. **D**, Structural comparison of di-nucleosome. The di-nucleosome from the Rpd3L–di-NCP complex was aligned to the ISW1-bound di-nucleosome structure (PDB ID: 7X3T) (56) using nucleosome 1 as a reference. Only the nucleosome cores are shown for comparison; associated proteins, including ISW1, are omitted. Histone octamers are shown in grey. DNA is shown in pink (Rpd3L–di-NCP) and blue (PDB ID: 7X3T). A pronounced bend in the linker DNA of Rpd3L–di-NCP induces an ∼90° rotation of nucleosome 2 relative to 7X3T. **E**, Enlarged view of the Sds3-Sin3^PAH3^ motif in the proximal module, showing its interaction between the Sds3-Sin3^PAH3^ motif of Rpd3L and the nucleosomal DNA at SHL+3.5 of the second nucleosome. **F**, Electrostatic surface potential representation of the Sds3-Sin3^PAH3^ motif, highlighting the formation of a basic pocket that engages with the acidic DNA backbone, shown in the same orientation as in E. **G**, Enlarged view of Sap30 interactions with the PAH3 and HID domains of Sin3 in the proximal module, with interacting residues involved in hydrogen bonding labeled(25,26,60). Hydrogen bonds are shown as blue dashed lines.

Strikingly, we observed additional densities at the linker DNA region between the two nucleosomes, which were absent in the mono-nucleosome structure. We employed AlphaFold3 predictions to generate candidate models and structural fitting finally revealed these densities correspond to the Sin3 PAH1 and PAH2 domains (Fig. 4B and C, Supplementary Fig. S11A). These domains contact the linker DNA between nucleosomes, with PAH1 forming long-range electrostatic contacts with the entry DNA of nucleosome 2 via conserved residues K232, K250, and K253 (Supplementary Fig. S11C). Consistently, truncation of the PAH1 and PAH2 domains abolished the preferential binding to di-nucleosomes (Supplementary Fig. S1B and S11D). Comparison with canonical di-nucleosome models revealed that the linker DNA adopts a markedly bent trajectory in our structure(56). We propose that this deformation is driven by PAH1/2-mediated interactions, which stabilize both the linker DNA conformation and the positioning of nucleosome 2 (Fig. 4D). Notably, the hydrophobic cleft of the PAH1 domain is fully exposed (Fig. 4C, Supplementary Fig. S11B), forming a conserved binding surface for potential transcriptional corepressors, and spatially allowing them for sequence-specific DNA engagement (57–59).

By reshaping the linker DNA trajectory through PAH1/2 domain engagement, nucleosome 2 is brought into proximity with the distal module of Rpd3L, where it becomes captured via weak electrostatic interactions by a basic pocket formed by Sds3 and the Sin3 PAH3 domain (Fig. 4E and F). Notably, the positioning of PAH3 in this context is dependent on its interaction with Sap30(60), enabling its engagement with nucleosome 2. In contrast, within the Rpd3S complex, the Sin3 PAH3 domain adopts a distinct conformation, rotated by approximately 180° relative to its orientation in Rpd3L (Fig. 4G, Supplementary Fig. S12). This observation indicates the structural adaptability of Sin3, whose multi-domain architecture enables alternative binding modes with specific subunits, resulting in distinct conformational states and clearly differentiated biological functions.

Together, these observations support a ‘guided landing’ architecture, in which the Sin3 PAH1/2 domains function as DNA-sensing elements that dock onto linker DNA only after stable engagement of nucleosome 1 by the proximal module. This sequential assembly serves to orient and stabilize nucleosome 2, positioning it for further capture by PAH3/Sds3. Importantly, the PAH1/2 domains also harbour a conserved hydrophobic cleft, potentially accommodating transcriptional corepressors with sequence-specific DNA-binding capacity. This two-step engagement model provides a mechanistic basis for Rpd3L’s di-nucleosome preference, highlighting a cooperative and topology-sensitive assembly strategy, in which Sin3 PAH domains are differentially deployed to interpret nucleosome spacing and guide hierarchical chromatin engagement.

### Molecular mechanism for the context-dependent deacetylation of Rpd3L

To biochemically explore the context-depended deacetylation by Rpd3L, we employed an endogenous octamer extraction approach to reconstitute both mono- and di-nucleosomes carrying native histone modifications. Using site-specific acetylation antibodies, we monitored Rpd3L-mediated deacetylation and quantified enzymatic efficiency by titrating Rpd3L concentrations against mono- or di-nucleosome substrates (Fig. 5A). Rpd3L effectively removed acetylation marks on both H3 and H4 tails across substrates, consistent with previous reports(61). Notably, using a sensitive H2BK12ac-specific antibody, we detected modest deacetylation of H2BK12ac. Moreover, high-throughput liquid chromatography coupled to tandem mass spectrometry (LC-MS/MS) further revealed reproducible deacetylation of H2AK5ac and H2AK9ac (Supplementary Fig. S13A). While previous studies have occasionally hinted at Sin3 HDAC activity beyond H3/H4, our findings provide comprehensive evidence that Rpd3L can target H2A and H2B tails in a chromatin context.

**Fig. 5.**
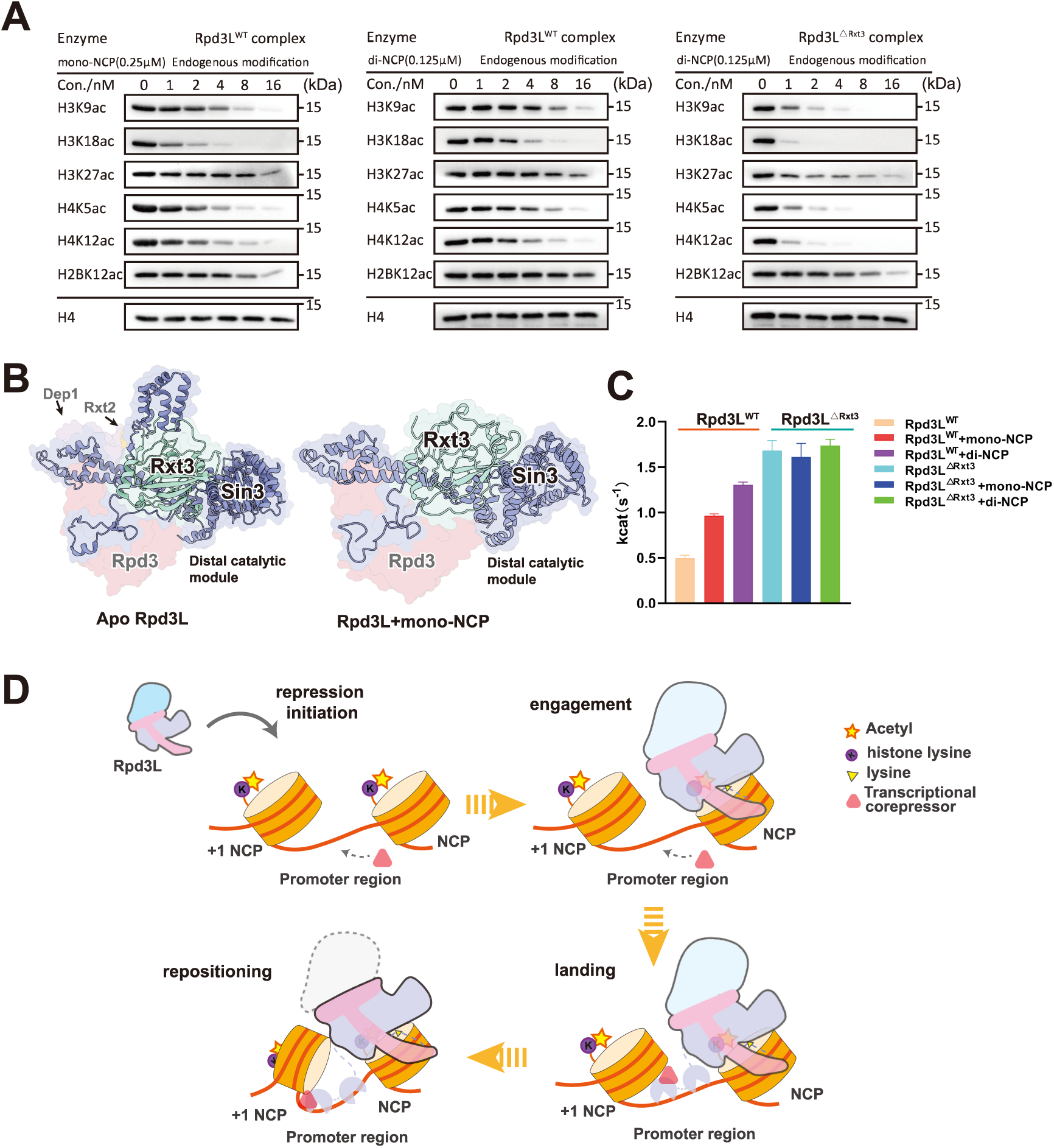
Substrate-induced activity regulation of the Rpd3L complex. **A**, Representative HDAC assay measuring Rpd3L activity on mono-nucleosomes (left) and di-nucleosomes (middle) reconstituted with endogenous histone octamers. Representative HDAC assay measuring Rpd3L^Δ1Rxt3^ activity on di-nucleosomes reconstituted with endogenous histone octamers (right). **B**, Structural comparison of the distal catalytic module in apo Rpd3L (PDB ID:8HPO) (left) versus mono-NCP-bound Rpd3L (right). The Rxt3 subunit engages tightly with components of the distal module upon substrate binding, contributing to its structural stabilization. **C**, HDAC activity of Rpd3L and Rpd3L^Δ1Rxt3^ measured using Boc-Lys(Ac)-AMC. Bar graph shows k_cat_ values for apo, mono-NCP, and di-NCP binding states. Nucleosome binding enhances Rpd3L activity in an Rxt3-dependent manner, with di-NCP inducing the strongest activation. Experiments were performed independently three times, and with error bars indicating the SD of the Kcat values (n = 3). **D**, Schematic model illustrating the molecular mechanism for the hierarchical deacetylation of Rpd3L.

To investigate the molecular basis of the two-step engagement-guided deacetylation, we examined conformational changes in Rpd3L upon binding to di-nucleosome substrates. The most notable difference was increased flexibility in the distal catalytic module (Fig. 4A, Supplementary Fig. S14). Thus, we hypothesized that this distal module flexibility might be essential for Rpd3L’s catalytic activation. Structural analysis of apo Rpd3L revealed that this module is constrained by interactions between Rxt3 and Sin3, suggesting a conformational gating mechanism (Fig. 5B). To test this, we generated an Rxt3 deletion mutant (Rpd3L^Δ1Rxt3^) that removed this constraint entirely and successfully purified Rpd3L^Δ1Rxt3^ as an intact complex (Supplementary Fig. S1B). This mutant exhibited elevated deacetylase activity in AMC-based assays, regardless of substrate context (apo, mono- or di-nucleosome, Fig. 5C), and showed enhanced activity toward di-nucleosomes in site-specific assays (Fig. 5A). Collectively, these findings suggest that chromatin-induced release of the distal module underlies Rpd3L activation, while Rxt3 serves as a regulatory subunit that suppresses catalytic output in the absence of appropriate chromatin engagement.

Given the above facts, we proposed the following working model (schematic Fig. 5D). Rpd3L operates through a hierarchical, chromatin-context-dependent mechanism to regulate its deacetylase activity. The process is initiated by the proximal catalytic module anchoring onto the first nucleosome, establishing a stable engagement with chromatin. Subsequently, the Sin3 PAH1/2 domains function as DNA-sensing modules that dock onto linker DNA, guiding the positioning of nucleosome 2 into proximity with the complex. This repositioning enables nucleosome 2 to be tethered by the PAH3/Sds3 interface, thereby inducing a structural rearrangement that unlocks the distal catalytic module. This allosteric activation transforms the distal site from a latent to an active state, enhancing Rpd3L’s catalytic breadth.

Together, this model illustrates how Rpd3L interprets chromatin topology through its asymmetric modules, integrating spatial chromatin cues into a finely tuned, stepwise enzymatic response. Such a mechanism enables locus-specific deacetylation and provides a framework for understanding how asymmetrical chromatin complexes achieve precision in chromatin regulation.

## Discussion

The activity of chromatin HDACs must be precisely regulated to preserve chromatin homeostasis. Through integrative cryo-EM and biochemical analyses, our study uncovers a hierarchical, chromatin-context-dependent mechanism by which the yeast Sin3 complex Rpd3L interprets nucleosomal substrates to regulate its deacetylase activity. Upon binding to mono- and di-nucleosome substrates, Rpd3L undergoes a substrate-dependent conformational rearrangement that allosterically activates its distal catalytic module. This activation is mediated by a reorganization of the coiled-coil scaffold, which propagates structural changes across the complex. Similar substrate-induced coiled-coil transitions have been reported in other regulators(62), suggesting a potentially conserved principle of chromatin-encoded allosteric control.

While Sin3 has long been regarded as a scaffolding platform, our structural findings highlight its active role in substrate sensing and complex activation. Specifically, we demonstrate that the Sin3 PAH1/2 domains engage linker DNA between nucleosomes, not only stabilizing Rpd3L’s binding to chromatin, but also positioning it in proximity to transcriptional corepressors frequently associated with these genomic regions. This suggests a dual function of PAH domains: as structural mediators for complex docking, and as potential recruitment hubs for sequence-specific regulators. This modular flexibility provides a mechanistic basis for how Sin3-containing HDAC complexes assemble with regulatory partners and adapt to diverse chromatin environments.

Notably, our structure of Rpd3L bound to a di-nucleosome substrate represents the highest-resolution model of a Sin3 HDAC complex engaging physiologically relevant chromatin. The topology-driven engagement and asymmetric activation of its two catalytic modules emphasize the importance of using native-like nucleosome templates to elucidate regulatory mechanisms. However, despite efforts, histone modification cues such as H3K4me3 recognition(63) was not observed in our studies, suggesting that either its recognition is transient, or that additional cofactors may be required to stabilize this interaction under physiological conditions.

Beyond static interactions, di-nucleosome engagement increases conformational flexibility and catalytic potential of the distal module, as supported by activity assays and structural analyses. These observations prompted us to investigate how nucleosome context shapes Rpd3L activity. We find that Rpd3L does not recognize acetyl-lysine residues solely by their identity or position; rather, nucleosome organization dictates the accessibility and efficiency of deacetylation. On di-nucleosomes, H4K5ac and H4K12ac—marks of newly deposited histones—are preferentially targeted, whereas on mono-nucleosomes, H3K18ac and H3K23ac are more efficiently deacetylated (Supplementary Fig. S13B and C). These results support a model in which Rpd3L fine-tunes its enzymatic specificity through substrate conformation and accessibility, rather than strict recognition of pre-existing acetylation marks.

Building on these insights, our model has important implications for understanding the physiological adaptability of Rpd3L. Chromatin architecture is dynamically remodelled in response to environmental signals such as nutrient availability, and pathogen invasion. These conditions require rapid transcriptional reprogramming, in part mediated by context-dependent modulation of histone acetylation. The asymmetric architecture of Rpd3L may provide a selective advantage in such scenarios. For instance, the phenomenon of Transcriptional Repression Memory (TREM) depends on the ability of chromatin regulators to remember and respond more efficiently to previously encountered stress(64,65). Rpd3L’s asymmetrical modular design, as well as the stepwise activated mechanism allows the complex to sense and respond to both local chromatin features and broader environmental signals, which offers a mechanistic speculation for such memory effects.

Taken together, our study reveals a previously unrecognized, hierarchical regulatory mechanism in which Rpd3L integrates spatial chromatin cues, structural dynamics, and environmental inputs to fine-tune its enzymatic activity. This asymmetric module-based regulation allows the complex to distinguish routine transcriptional repression from more complicated events. By enabling both immediate and reinforced repression responses, Rpd3L’s two-module architecture transforms transient chromatin interactions into a durable and adaptive transcriptional memory. This mechanistic framework may represent a general strategy by which chromatin effectors integrate structural and epigenetic information to guide gene regulatory decisions in eukaryotic cells.

## Data Availability

The cryo-EM density maps and corresponding atomic coordinates have been deposited in the Electron Microscopy Data Bank (EMDB) and Protein Data Bank (PDB) under accession numbers EMD-64741 and PDB-9V2V for the Rpd3L-mono-NCP complex; EMD-64742 and PDB-9V2W for the Rpd3L-di-NCP complex.

## Acknowledgements

We thank the cryo-EM Center, Guangzhou Institutes of Biomedicine and Health Chinese Academy of Sciences and Advanced Bio-imaging Technology Platform of Guangzhou Laboratory for cryo-EM beamtime and all staff members for their assistance with data collection. We also thank the Mass Spectrometry System staff members at the National Facility for Protein Science in Shanghai (NFPS), Zhangjiang Lab, China for technical support and assistance in XL-MS data collection and analysis.

## Author Contributions

J.H. and H.L. designed the project. H.L. and X.Y. constructed the expression plasmids and prepared protein complexes involved in the research. H.Z., C.W., N.Z. and Y.Z. prepared the nucleosomes involved in the research and reconstituted protein-nucleosome complexes for the cryo-EM studies. H.L. and C.W. performed biochemical assays. H.Z., H.L. and J.H. conducted cryo-EM experiments and determined the structures. H.Z, H.L. S.D. and B.Z. prepared the structures and figures. L.Y. and Y.X. performed the quantitative LC-MS/MS experiment and analysed the data. Y.Z., W.C., D.Q. and D.P. discussed the results and provided feedback in the manuscript. H.Z. and J.H. prepared the manuscript with input from all authors. J.H. supervised and directed the overall research.

