## Supplemental Data 1 for "Chromatin context-dependent deacetylation by the asymmetric Rpd3L"

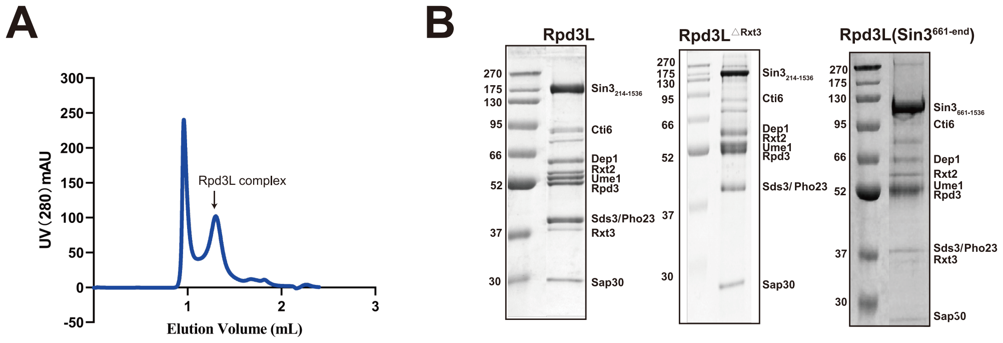

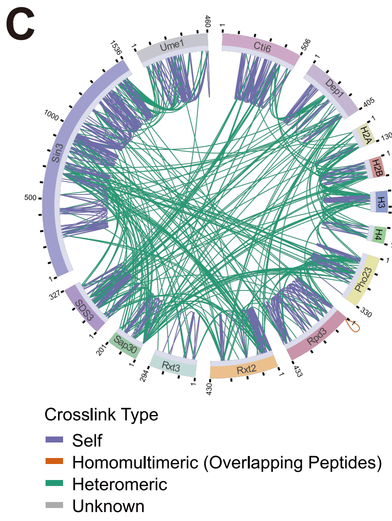

E

| protein1(site)-protein2(site) | #Spectrum Number | #Best E-value |
|---|---|---|
| H2A(37)-H2B(83) | 17 | 1.36E-14 |
| H2B(106)-SDS3(199) | 13 | 7.69E-08 |
| H2A(120)-H3(37) | 11 | 1.34E-13 |
| H3(80)-H4(78) | 10 | 5.88E-24 |
| H2A(37)-H4(92) | 9 | 1.46E-12 |
| Cti6(383)-H2B(118) | 9 | 1.97E-08 |
| H3(28)-Pho23(154) | 6 | 4.85E-11 |
| H2B(32)-H4(92) | 5 | 8.98E-12 |
| H2B(106)-Sap30(2) | 5 | 9.64E-09 |
| H3(28)-Sap30(69) | 3 | 4.50E-06 |
| H3(19)-Rpd3(98) | 3 | 4.37E-11 |
| Cti6(169)-H2B(117) | 3 | 5.64E-10 |
| Cti6(383)-H2A(96) | 3 | 3.64E-05 |
| H3(57)-SDS3(199) | 2 | 2.27E-07 |
| Cti6(382)-H3(19) | 2 | 2.43E-04 |
| Dep1(79)-H2A(102) | 2 | 7.29E-08 |

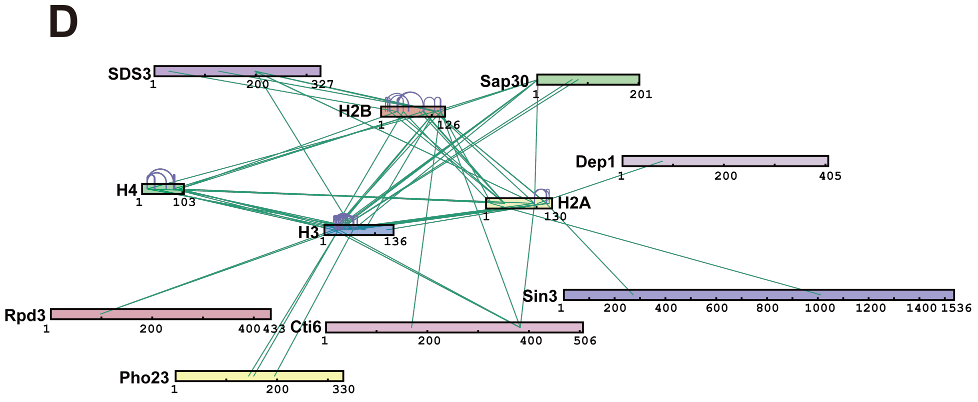

**Supplementary Figure S1. Purification of the Rpd3L complex and XL-MS analysis of Rpd3L-mono-NCP interaction.**

**A**, Representative size-exclusion chromatography profile of purified Rpd3L complex. **B**, SDS-PAGE analysis of purified Rpd3L complexes, including the apo Rpd3L complex, the Rxt3-deficient complex (Rpd3L$^{\Delta Rxt3}$), and an Rpd3L complex containing a Sin3 construct lacking the PAH1 and PAH2 domains (Rpd3L(Sin3$^{661\text{-end}}$)). The presence of the expected subunits is indicated. **C**, Circular plot of inter-subunit cross-links identified by XL-MS for Rpd3L-mono-NCP interaction. **D**, Focused XL-MS mapping of cross-links involving the nucleosomal histone octamer. **E**, Representative cross-linked sites between Rpd3L subunits and histones identified by cross-linking mass spectrometry (XL-MS) of the Rpd3L-mono-NCP are listed.

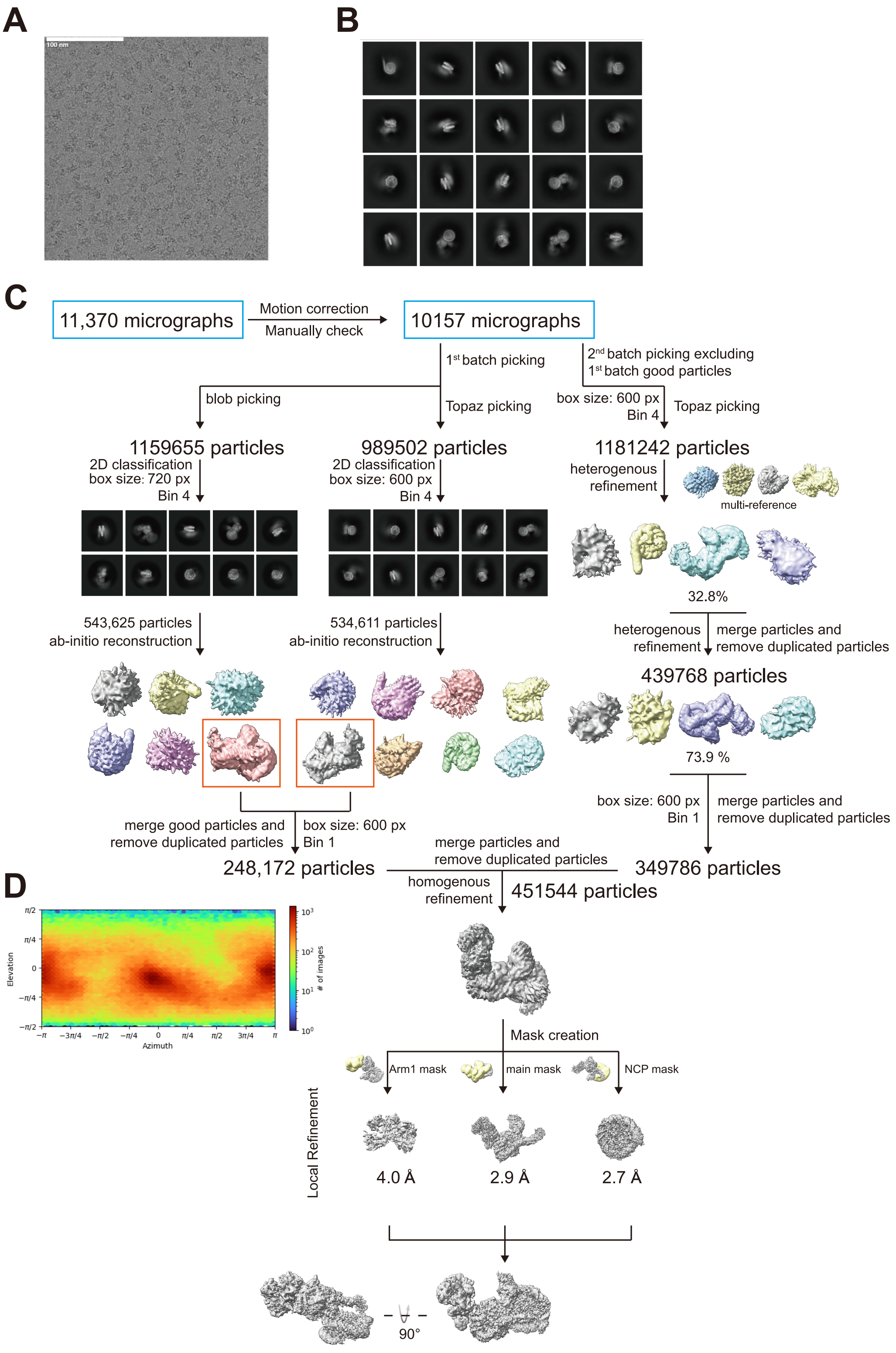

**Supplementary Figure S2. Cryo-EM data processing pipelines for the Rpd3L-mono-NCP complex datasets.**

Representative cryo-EM image(**A**), 2D classification(**B**) and flow-charts(**C**) of the cryo-EM images processing and 3D reconstruction for Rpd3L-mono-NCP in cryoSPARC. Rpd3L-mono-NCP was further segmented into three regions: the distal catalytic module, the main body, and the nucleosome. Each region was individually masked for particle subtraction and local refinement in cryoSPARC. The resulting focused maps were then merged into a single composite map using ChimeraX. **D**, Particle angular distribution of Rpd3L-mono-NCP.

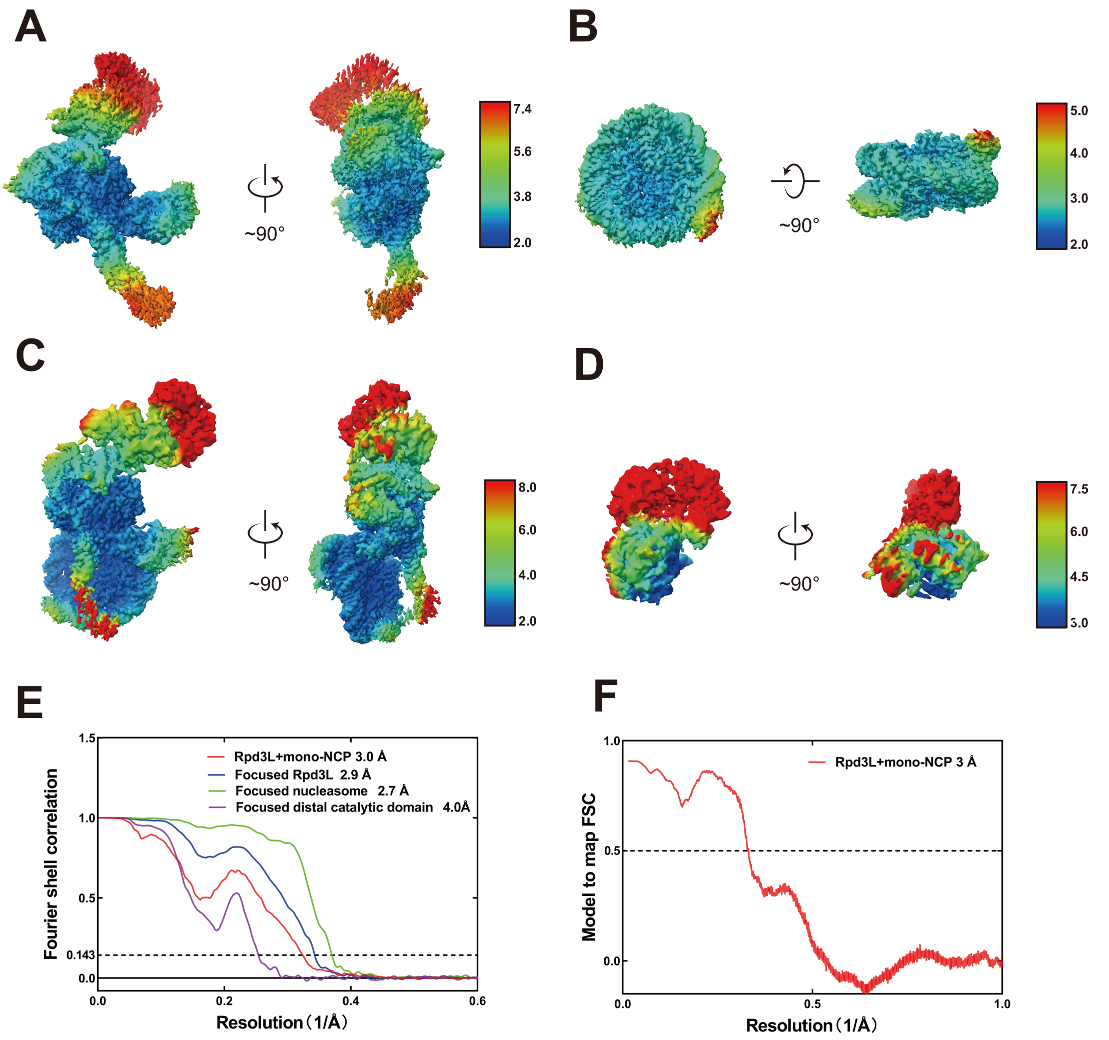

**Supplementary Figure S3. Resolution assessment of the Rpd3L-mono-NCP cryo-EM structures.**

**A-D**, Local resolution assessments for the focused Rpd3L (A), focused NCP (B), Rpd3L bound to mono-NCP (C) and focused distal catalytic module structures (D). **E**, Global resolution assessment by Fourier shell correlation (FSC) at the 0.143 criterion. **F**, Model to map FSC at the 0.5 criterion.

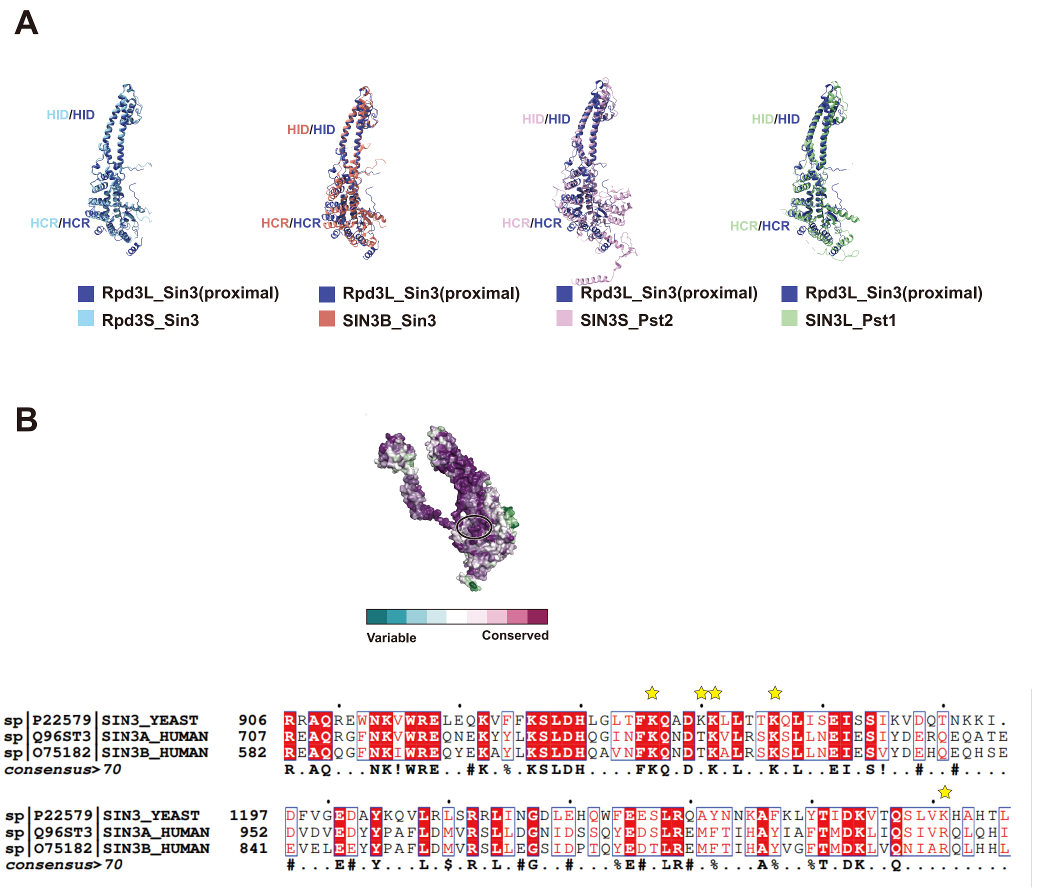

**Supplementary Figure S4. Conserved features of the Sin3 subunit in Rpd3L and its homologs across species.**

**A**, Structural alignment of the Sin3 subunit from *S. cerevisiae* Rpd3L (dark blue) with Sin3 homologs including *S. cerevisiae* Rpd3S (sky blue), human SIN3B (orange), *S. pombe* SIN3S (pink), and *S. pombe* SIN3L (green), using the HID (HDAC interaction domain) as a reference， reveals that the HCR (highly conserved region) is structurally well conserved across species. **B**, ConSurf analysis and multiple sequence alignment of *S. cerevisiae* Sin3 with human SIN3A and SIN3B highlight conserved regions and nucleosome-binding interfaces. Conserved residues are shaded from cyan (variable) to purple (highly conserved). Black ovals indicate nucleosome-binding regions; key interface residues are marked with yellow asterisk.

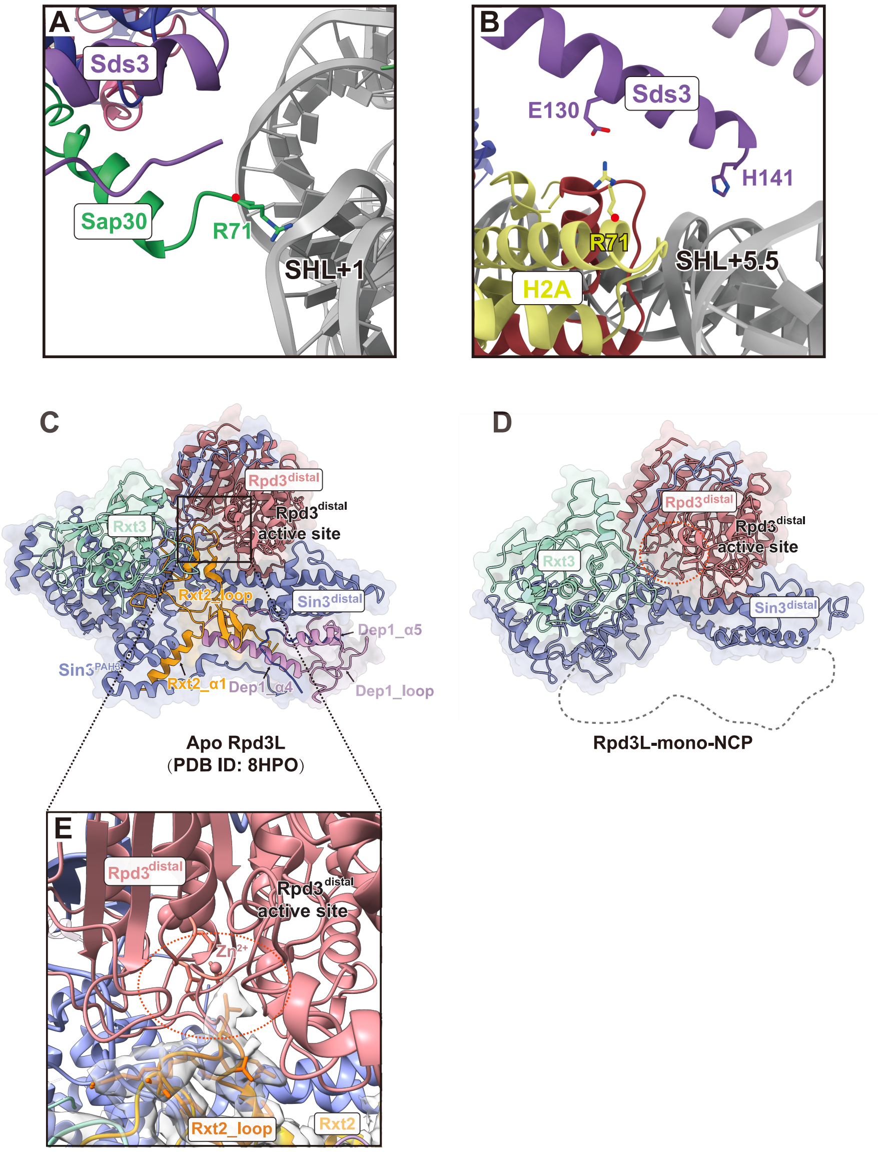

**Supplementary Figure S5. Structural focus on Rpd3L–mono-NCP interface and distal catalytic module.**

**A**, Enlarged view of SAP30 interacting with nucleosomal DNA at SHL+1. **B**, Enlarged view of Sds3 interacting with nucleosomal DNA at SHL +5.5 and with the H2A histone. **C**, Structure of the distal catalytic module of apo Rpd3L (PDB ID: 8HPO), highlighting the interface between the Rxt2 loop and the Rpd3 active site. **D**, Distal catalytic module of Rpd3L in the mono-nucleosome – bound state, with regions that become flexible upon nucleosome binding indicated by dashed circles. **E**, Close-up view of the interface between the Rxt2 loop and the Rpd3 active site in apo Rpd3L. The cryo-EM density of the Rxt2 loop is shown as a transparent surface, while the corresponding atomic model is highlighted in dark orange with side chains displayed. The Rpd3 active site is indicated by an orange dashed circle.

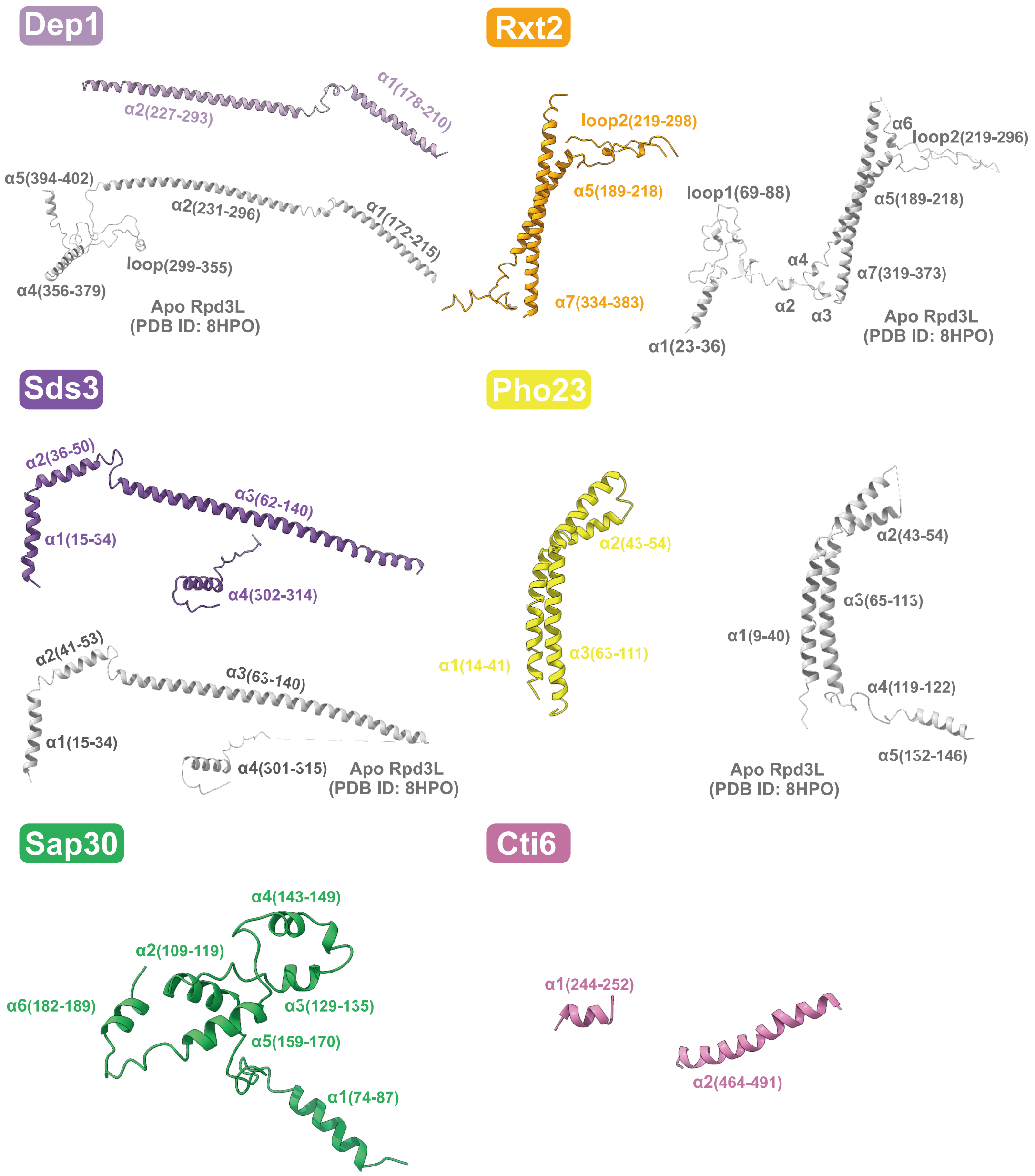

**Supplementary Figure S6. Structures of Rpd3L subunits in apo and mono-NCP bound states.**

Dep1, Sds3, Rxt2, Pho23, Sap30 and Cti6 are shown in both the apo state (grey) and the Rpd3L-mono-NCP bound state (coloured), highlighting their structural features under each condition.

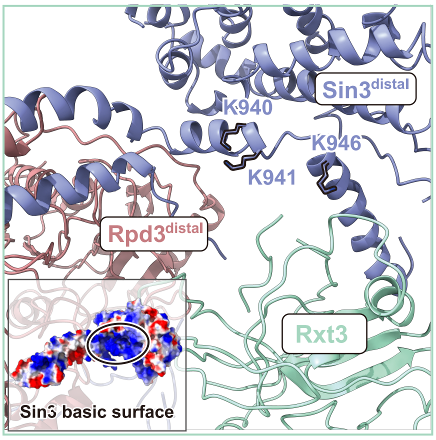

**Supplementary Figure S7. Rxt3-mediated occlusion of the NCP-binding surface on Sin3$^{distal}$.**

Steric occlusion of the DNA-binding surface of Sin3$^{distal}$ by Rxt3. The electrostatic surface potential (-/+5.0 kT/e) is shown on the Sin3$^{distal}$ basic surface.

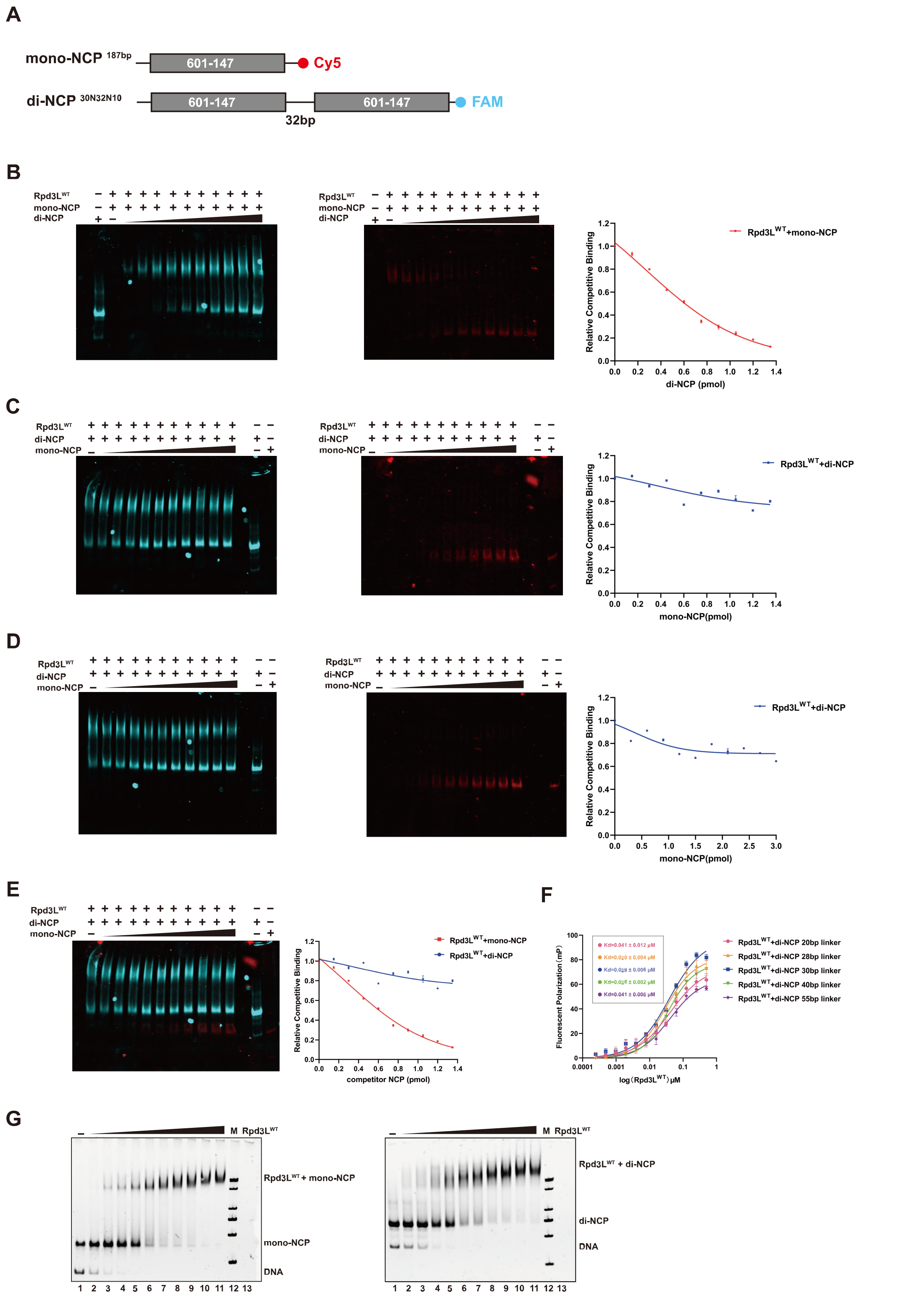

**Supplementary Figure S8. Competitive binding assays reveal higher affinity of Rpd3L for di-nucleosomes.**

**A**, Fluorescent labeling schematic of mono-NCP and di-NCP. **B**, Fluorescent EMSA showing competition between di-NCP and mono-NCP. Increasing concentrations of di-NCP reduced mono-NCP binding, indicating higher affinity of Rpd3L for di-NCP. Band intensities of Rpd3L-mono-NCP were quantified based on grayscale values and plotted on the right. **C**, Reverse competition assay where increasing mono-NCP concentrations were used to displace Rpd3L from di-NCP. Di-NCP binding remained largely unchanged. Band intensities of Rpd3L-di-NCP were quantified based on grayscale values and plotted on the right. **D**, Competition assay with a two-fold increased gradient of mono-NCP. Rpd3L retained preferential binding to di-NCP, despite elevated mono-NCP levels. Band intensities of Rpd3L-di-NCP were quantified based on grayscale values and plotted on the right. **E**, merged fluorescent image from b, c and quantification of signal intensity showing that di-NCP effectively outcompetes mono-NCP for Rpd3L binding. All fluorescence intensity (grayscale) was measured in triplicate for each condition, and the values represent the average of these three measurements. **F**, Fluorescence polarization analysis of Rpd3L binding to di-NCP with varying linker DNA lengths. Each measurement was performed in triplicate. The fitted Kd values are indicated, with the standard error (SE) calculated from the curve fitting procedure. **G**, EMSA assays showing Rpd3L binding to mono-NCP (left) and di-NCP(right).The amount of nucleosome in each reaction was 1 pmol, and Rpd3L was added at increasing concentrations of 0, 0.3, 0.6, 0.9, 1.2, 1.5, 1.8, 2.1, 2.4, 2.7, and 3.0 pmol.

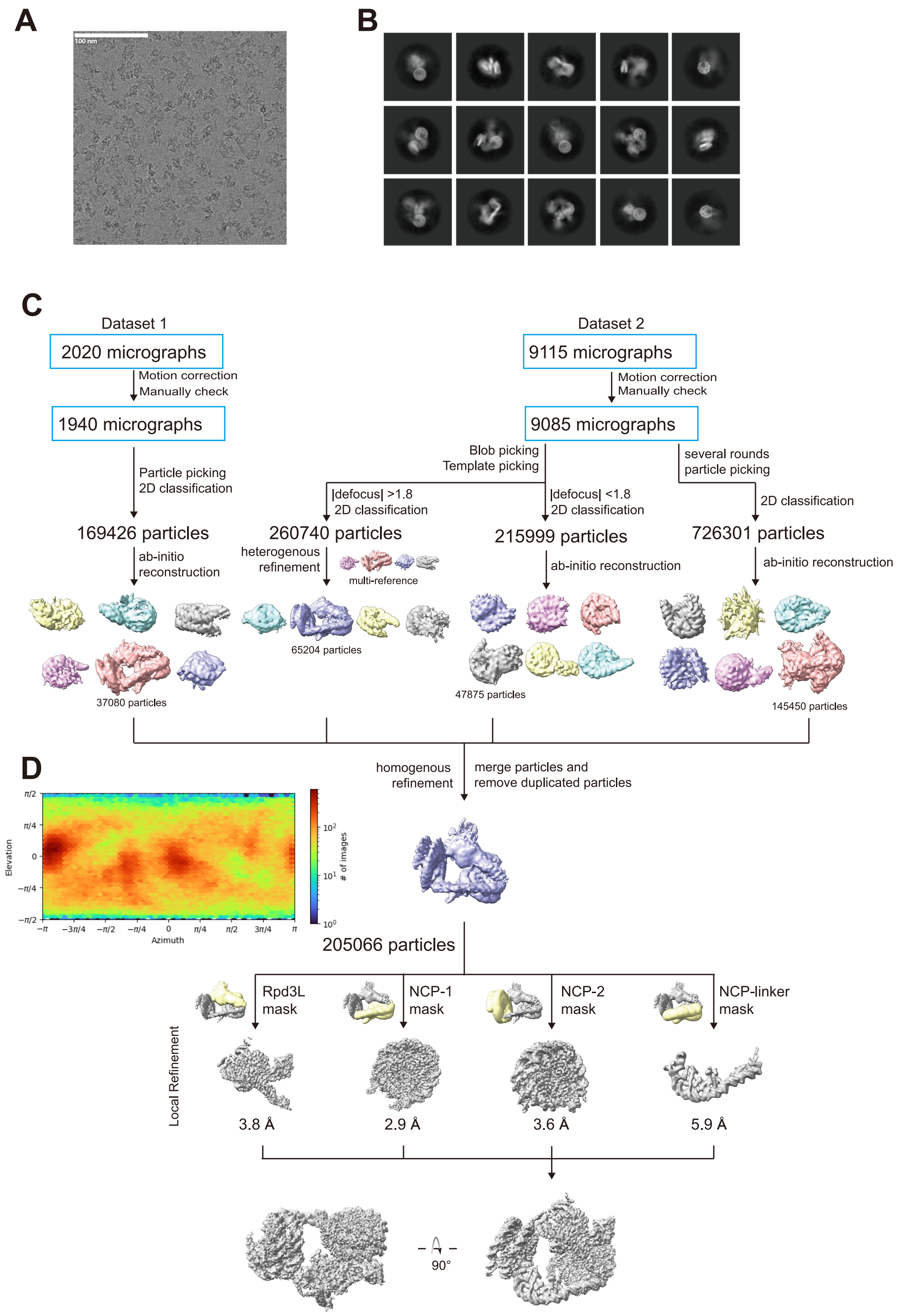

**Supplementary Figure S9. Cryo-EM data processing pipelines for the Rpd3L-di-NCP complex datasets.**

**A**, Representative cryo-EM micrograph. **B**, Representative 2D class averages. **C**, Flowchart of cryo-EM data processing and 3D reconstruction in cryoSPARC. The complex was divided into four regions: Rpd3L, DNA linker, and two nucleosomes for particle subtraction and local refinement. Focused maps were combined using ChimeraX. **D**, Particle angular distribution of Rpd3L-di-NCP.

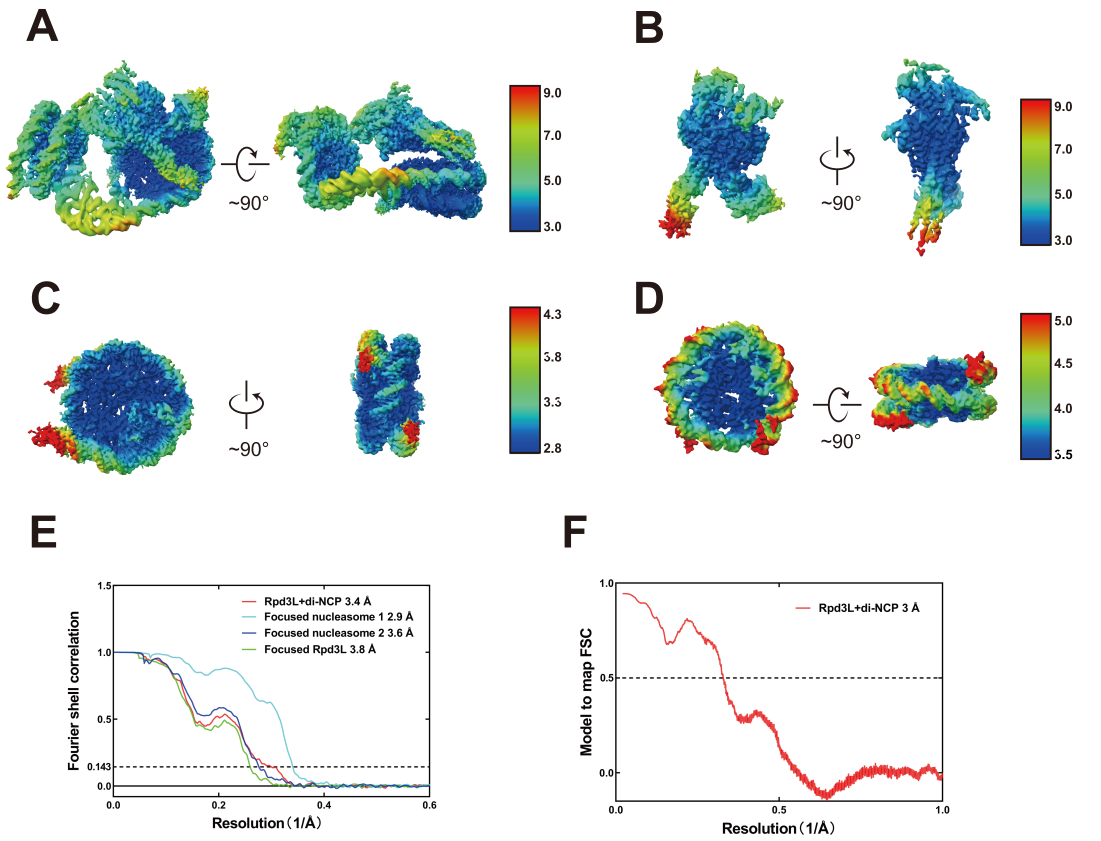

**Supplementary Figure S10. Resolution assessment of the Rpd3L-di-NCP cryo-EM structures.**

**A-D**, Local resolution assessments for the Rpd3L-di-NCP structures(A)， focused Rpd3L (B) , focused nucleasome 1 (C) and nucleasome 2 (D). **E**, Global resolution assessment by Fourier shell correlation (FSC) at the 0.143 criterion. **F**, Model to map FSC at the 0.5 criterion.

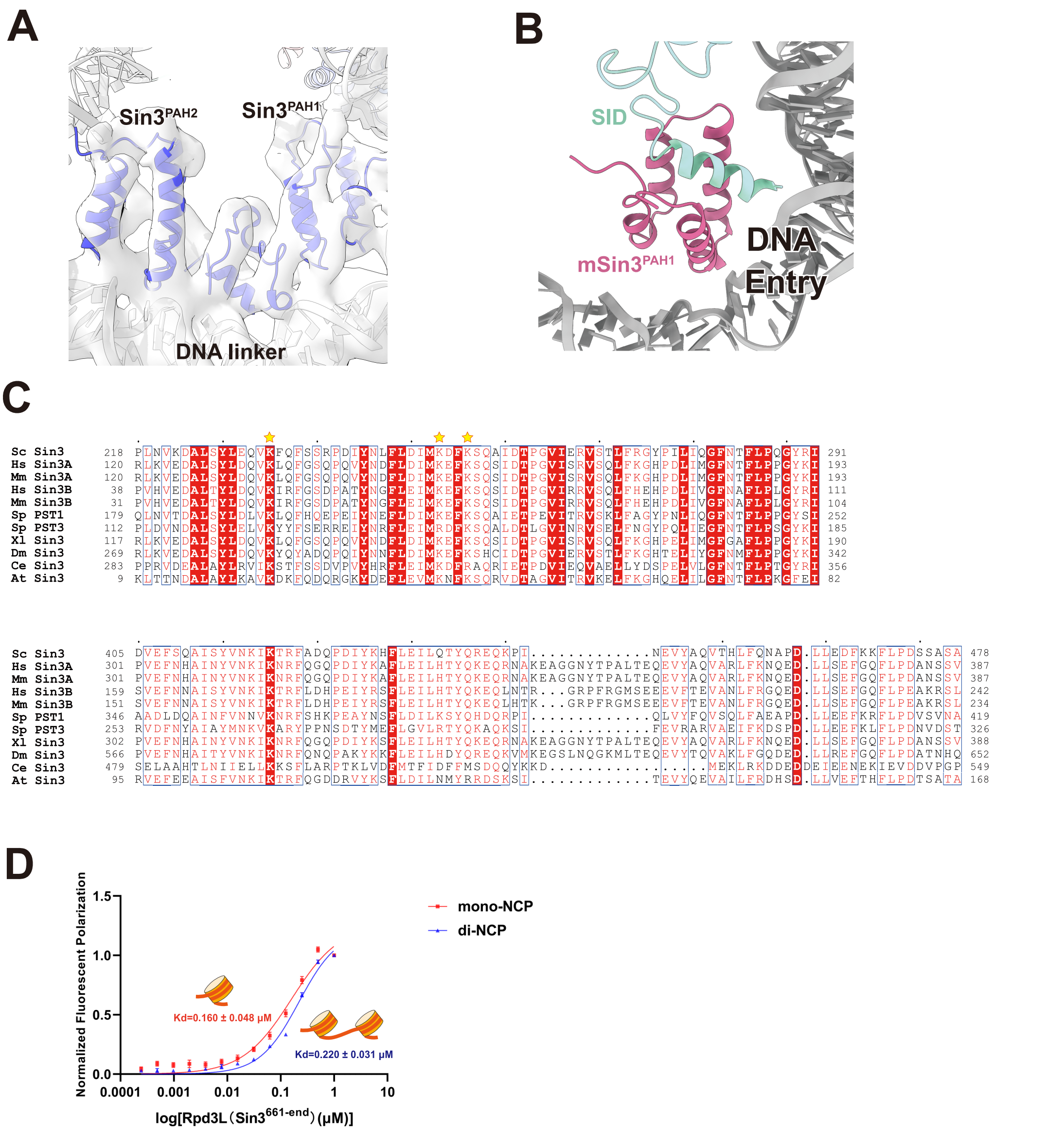

**Supplementary Figure S11. Structural and evolutionary conservation of Sin3 PAH domains.**

**A**, Structural view of the PAH1 and PAH2 domains of the Sin3 subunit in the proximal catalytic region of the Rpd3L di-nucleosome complex, shown in the context of their association with linker DNA. The atomic model is fitted into the cryo-EM density, which is displayed as a transparent surface. **B**, The proposed binding model of PAH1 domain of Sin3 with transcription factors and the promoter region DNA linker, using Sap25-mSin3 PAH1 complex (PDB ID: 2RMS) as an example. The position of Sap25-mSin3 PAH1 is aligned with the PAH1 structure in Rpd3L-di-NCP. **C**, Sequence alignment of

Sin3$^{PAH1}$ and Sin3$^{PAH2}$ domains across species. Multiple sequence alignment of PAH1 (top) and PAH2 (bottom) domains from various organisms, including *Mus musculus* (Mm), *Homo sapiens* (Hs), *Xenopus laevis* (Xl), *Drosophila melanogaster* (Dm), *Schizosaccharomyces pombe* (Sp), *Saccharomyces cerevisiae* (Sc), *Arabidopsis thaliana* (At), and *Caenorhabditis elegans* (Ce). Basic residues that interact with DNA in the structure are marked with yellow asterisks above the alignment. **D**, Fluorescence polarization assay comparing binding of an Rpd3L complex containing a Sin3 construct lacking the PAH1 and PAH2 domains (Rpd3L(Sin3$^{661\text{-}end}$)) to mono-NCP and di-NCP substrates. Each measurement was performed in triplicate, with error bars showing the standard deviation (SD). The fitted Kd values are indicated, with the standard error (SE) calculated from the curve fitting procedure.

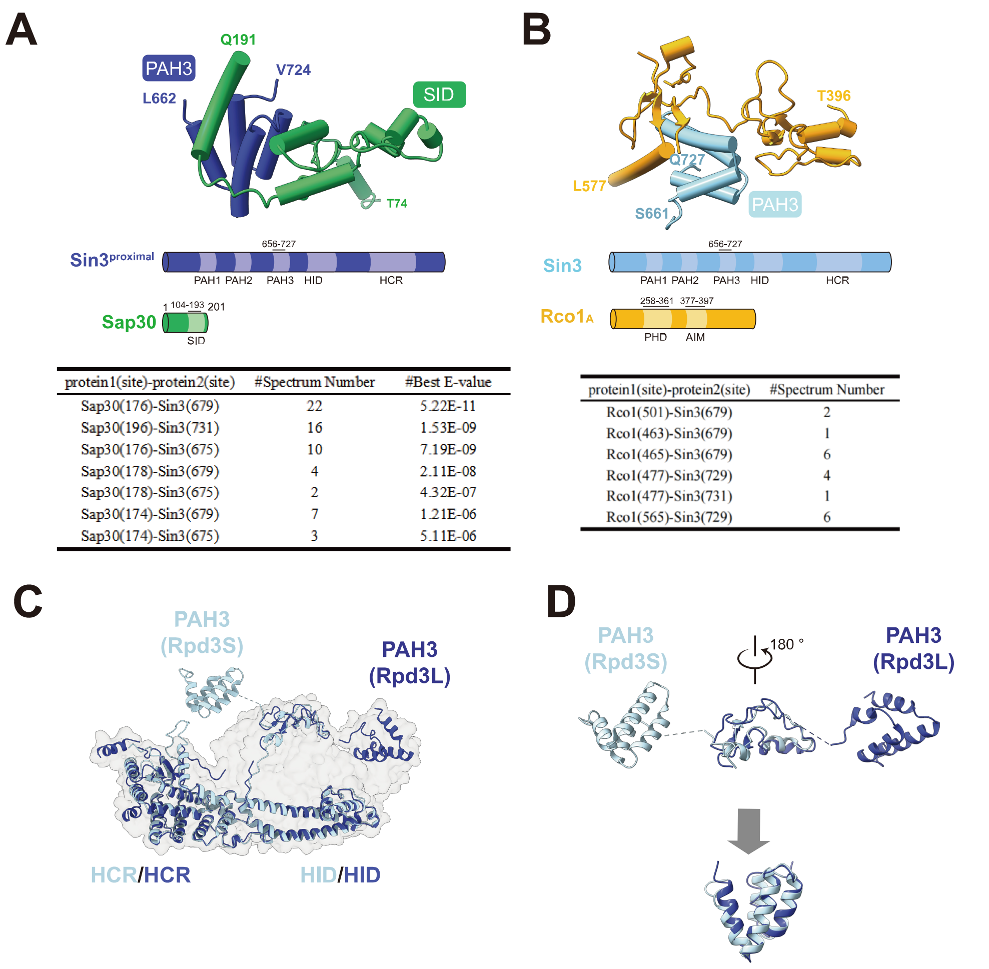

| protein1(site)-protein2(site) | #Spectrum Number | #Best E-value |
|---|---|---|
| Sap30(176)-Sin3(679) | 22 | 5.22E-11 |
| Sap30(196)-Sin3(731) | 16 | 1.53E-09 |
| Sap30(176)-Sin3(675) | 10 | 7.19E-09 |
| Sap30(178)-Sin3(679) | 4 | 2.11E-08 |
| Sap30(178)-Sin3(675) | 2 | 4.32E-07 |
| Sap30(174)-Sin3(679) | 7 | 1.21E-06 |
| Sap30(174)-Sin3(675) | 3 | 5.11E-06 |

| protein1(site)-protein2(site) | #Spectrum Number |
|---|---|
| Rco1(501)-Sin3(679) | 2 |
| Rco1(463)-Sin3(679) | 1 |
| Rco1(465)-Sin3(679) | 6 |
| Rco1(477)-Sin3(729) | 4 |
| Rco1(477)-Sin3(731) | 1 |
| Rco1(565)-Sin3(729) | 6 |

**Supplementary Figure S12. Structural comparison of Sin3 PAH3 domain interactions in Rpd3L and Rpd3S.**

**A**, Cryo-EM structure of the $Sin3^{PAH3}$ domain bound to Sap30 in Rpd3L, supported by crosslinking mass spectrometry analysis. **B**, Cryo-EM structure of the $Sin3^{PAH3}$ domain in the Rpd3S complex (PDB ID: 8KC7) bound to $Rco1_A$, also supported by crosslinking data. **C**, Structural alignment of Sin3 from Rpd3L (dark blue) and Rpd3S (light blue) using the HID domain as a reference, revealing an angular displacement of the PAH3 domain. **D**, Superposition of $Sin3^{PAH3}$ domains from Rpd3L and Rpd3S after a 180° rotation highlights conformational rearrangement enabling alignment.

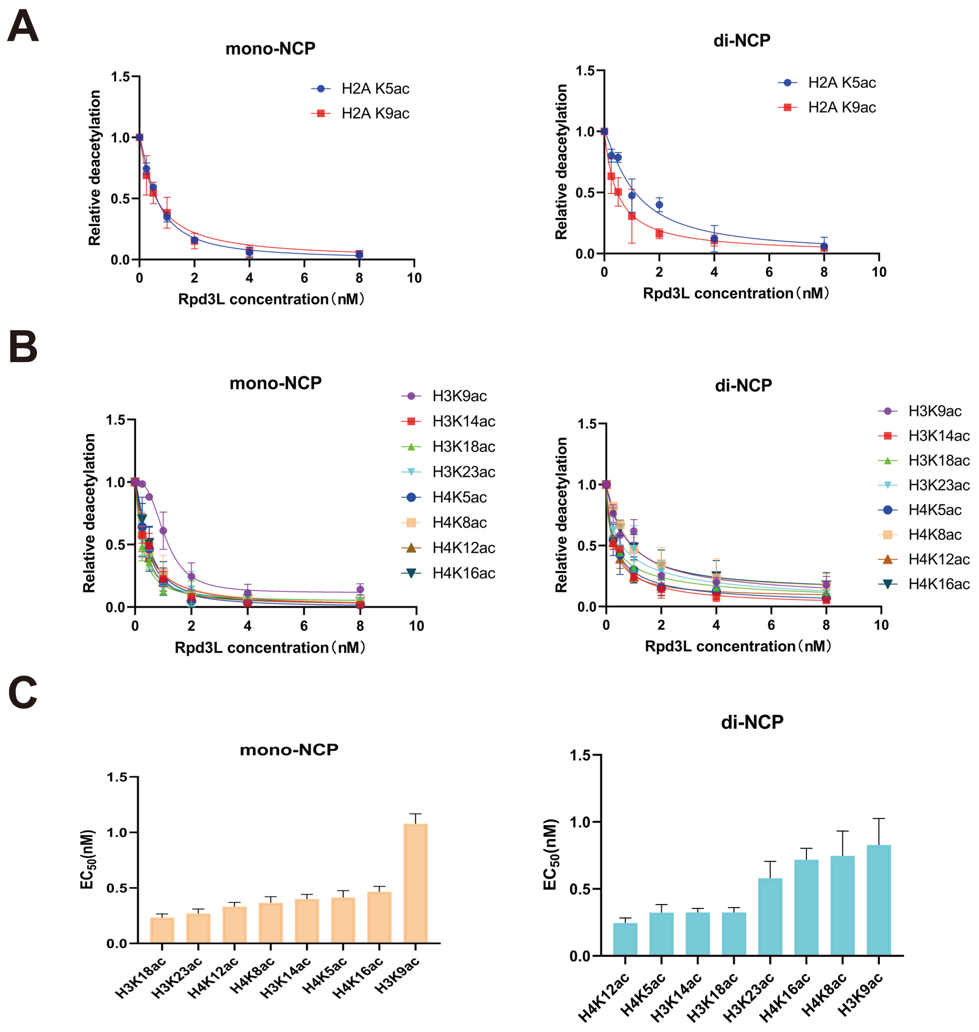

**Supplementary Figure S13. Quantitative mass spectrometry analysis of Rpd3L-mediated histone deacetylation.**

**A**, Representative deacetylation sites on histone H2A identified by quantitative mass spectrometry analysis of Rpd3L activity on mono-nucleosomes (left) and di-nucleosomes (right) reconstituted with endogenous histone octamers. **B**, Representative deacetylation sites on histone H3 and H4 identified by quantitative mass spectrometry analysis of Rpd3L activity on mono-nucleosomes (left) and di-nucleosomes (right) reconstituted with endogenous histone octamers. **C**, $EC_{50}$ values of Rpd3L deacetylation activity at representative lysine residues on histones H3 and H4, determined from the fitted curves shown in panel B. Measurements were performed using mono-nucleosome (left) and di-nucleosome (right) substrates. Higher

$EC_{50}$ values reflect lower enzymatic activity. Each experiment was performed in triplicate, and the values represent the average of the three measurements. Error bars represent the standard deviation (SD) of three independent experiments.

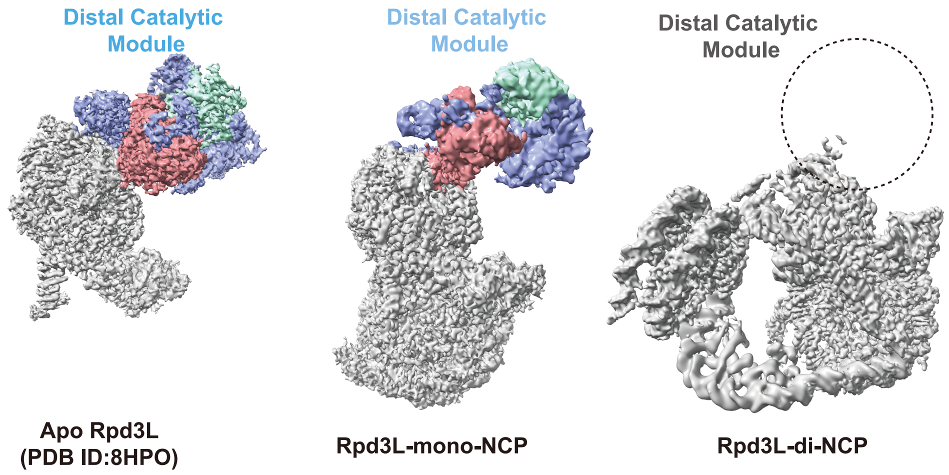

**Supplementary Figure S14. Structural comparison of apo Rpd3L, Rpd3L-mono-NCP and Rpd3L-di-NCP complexes.**

Cryo-EM maps of apo Rpd3L (left), Rpd3L bound to mono-NCP (middle), and Rpd3L bound to di-NCP (right) are shown. The distal catalytic module is highlighted in colour, while the rest of the complex is shown in grey. Increased flexibility of the distal catalytic module is observed upon nucleosome binding. In the Rpd3L-di-NCP complex, the distal catalytic module is not well resolved and is indicated by a dashed circle marking its expected position.

**Table. 1 | Cryo-EM data collection, refinement and validation statistics**

| | Rpd3L-mono-NCP (EMDB-64741) (PDB 9V2V) | Rpd3L-di-NCP (EMDB-64742) (PDB 9V2W) |
|---|---|---|
| **Data collection and processing** | | |
| Magnification | 165000 | 165000 |
| Voltage (kV) | 300 | 300 |
| Electron exposure (e–/Å$^2$) | 50 | 50 |
| Defocus range (μm) | 0.8-2.4 | 0.8-2.4 |
| Pixel size (Å) | 0.71 | 0.71 |
| Symmetry imposed | *C1* | *C1* |
| Initial particle images (no.) | 1159655 | 646165 |
| Final particle images (no.) | 451544 | 205066 |
| Map resolution (Å) | 3.0 | 3.4 |
| FSC threshold | 0.143 | 0.143 |
| Map resolution range (Å) | 3.02-24.92 | 3.01-13.64 |
| **Refinement** | | |
| Initial model used (PDB code) | PDB 8HPO/4LD9 | PDB 8HPO/4LD9 |
| Model resolution (Å) | 3.0 | 3.0 |
| FSC threshold | 0.5 | 0.5 |
| Map sharpening *B* factor (Å$^2$) | -32.95 | -36.66 |
| Model composition | | |
| Non-hydrogen atoms | 34241 | 39938 |
| Protein residues | 3411 | 3253 |
| Nucleotide | 310 | 658 |
| *B* factors (Å$^2$) | | |
| Protein | 134.72 | 83.43 |
| Nucleotide | 71.37 | 80.79 |
| R.m.s. deviations | | |
| Bond lengths (Å) | 0.017 | 0.010 |
| Bond angles (°) | 1.325 | 1.058 |
| Validation | | |
| MolProbity score | 1.93 | 1.87 |
| Clashscore | 8.88 | 10.13 |
| Poor rotamers (%) | 0.45 | 0.84 |
| Ramachandran plot | | |
| Favored (%) | 92.92 | 95.03 |
| Allowed (%) | 6.63 | 4.81 |
| Disallowed (%) | 0.95 | 0.16 |
